# Improved synthetic lipidation-based protein translocation system for SNAP-tag fusion proteins

**DOI:** 10.1101/2020.04.09.035188

**Authors:** Tatsuyuki Yoshii, Kai Tahara, Sachio Suzuki, Yuka Hatano, Keiko Kuwata, Shinya Tsukiji

## Abstract

The ability to artificially attach lipids to specific intracellular protein targets would be a valuable approach for controlling protein localization and function in cells. We recently devised a chemogenetic method in which a SNAP-tag fusion protein can be translocated from the cytoplasm to the plasma membrane by post-translationally and covalently conjugating a synthetic lipopeptide in cells. However, the first-generation system lacked general applicability. Herein, we present an improved synthetic lipidation system that enables efficient plasma membrane translocation of SNAP-tag fusion proteins in cells. This second-generation system is now applicable to the control of various cell-signaling molecules, offering a new and useful research tool in chemical biology and synthetic biology.

## MAIN TEXT

Protein lipidation involves the covalent modification of lipids to proteins, and is a central mechanism for localizing proteins on the surface of organelle membranes in cells.^1–4^ Cells regulate the subcellular localization of diverse proteins via lipidation, and lipidated (membrane-anchored) proteins play critical roles in various biological processes and cell signaling. In cells, protein lipidation is dictated by specific signal sequences. For example, the N-terminal MGXXXS/T sequence codes for myristoylation,^1,2^ and the C-terminal CAAX motif codes for prenylation.^4^ By fusing a lipidation signal sequence to a protein of interest, we can express the protein in lipidated form and enable protein targeting to a specific organelle surface. However, such genetic engineering approaches lack temporal control over the protein lipidation step. Because the localization and activity of many cellular proteins are drastically modulated by protein lipidation, the ability to post-translationally and chemically attach a lipid (or lipid analog) to a specific protein target would serve as a powerful chemical biology approach for the rapid and effective control of protein function in cells. Despite this potential, few attempts have been made to manipulate proteins in cells via such synthetic post-translational lipidation.^5^ Here, we present a synthetic post-translational protein lipidation system for use as a general tool to control protein localization and cell signaling in living cells.

The self-localizing ligand-induced protein translocation (SLIPT) technology is an emerging platform that enables the control of intracellular protein localization using synthetic self-localizing ligands (SLs).^6–9^ Based on the SNAP-tag labeling technique,^10^ we previously developed a SLIPT system targeting the inner leaflet of the plasma membrane (PM) using SNAP_f_,^11^ a fast-labeling mutant of the SNAP-tag protein (hereinafter referred to as SNAP) (**Fig. 1a**).^8^ In this system, an engineered SNAP_f_ containing a six-repeat of lysine residues (K6) at its N-terminus (K6-SNAP_f_) is used as a protein tag for fusion with a protein of interest. In addition, mgcBCP (**Fig. 1b**), a SNAP substrate ligand benzylchloropyrimidine (BCP) linked via flexible linker to a myristoyl-Gly-Cys (myrGC) lipopeptide motif,^12,13^ is used as an SL for K6-SNAP_f_. When mgcBCP is added to a culture medium of cells expressing a K6-SNAP_f_-fused protein of interest, mgcBCP enters the cells and relocates the protein to the PM as a result of the covalent attachment of the lipopeptidic myrGC motif to the SNAP_f_ domain (**Fig. 1a**).^8^ Therefore, the mgcBCP/K6-SNAP_f_ SLIPT system can be regarded as a synthetic post-translational protein lipidation system.

**Fig. 1.**
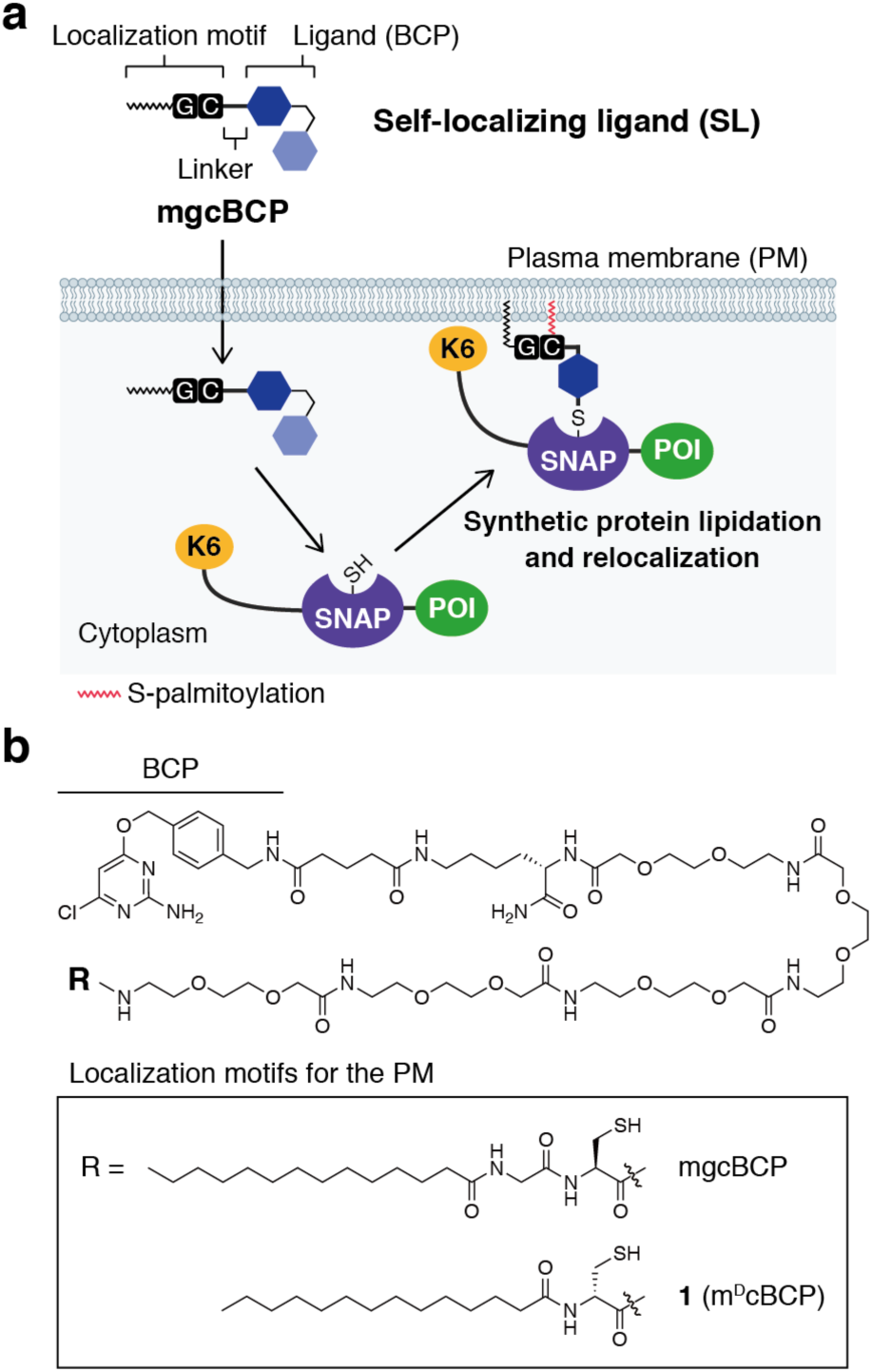
The SNAP SLIPT system based on synthetic post-translational protein lipidation. (**a**) Schematic illustration of the mgcBCP-based K6-SNAP SLIPT system (POI, protein of interest). (**b**) Chemical structures of SLs used in this study.

We then demonstrated the application of the lipidation-based K6-SNAP_f_ SLIPT system to induce the PM translocation of a guanine nucleotide exchange factor for Ras (Ras GEF), enabling synthetic activation of the endogenous Ras/ERK pathway.^8^ However, the original mgcBCP/K6-SNAP_f_ SLIPT system was not versatile enough for use as a general tool. For example, when we fused HaloTag as a fluorescent labeling tag to the C-terminus of K6-SNAP_f_ (K6-SNAP_f_-Halo), the K6-SNAP_f_-Halo construct showed only marginal PM localization by mgcBCP. Furthermore, we encountered difficulties in applying the original system to control other signaling proteins such as Tiam1. Therefore, we aimed herein to develop an improved SNAP SLIPT system by redesigning the SL and K6-SNAP_f_ construct.

During our previous work using a myrGC-tethered trimethoprim (mgcTMP), an SL for *E. coli* dihydrofolate reductase (eDHFR),^6^ we unexpectedly found that the myrGC motif undergoes degradation in cells (probably by proteases).^9^ To overcome this problem, we developed a designer myristoyl-^D^Cys (m^D^c) lipidic structure as a novel protease-resistant localization motif.^9^ We hypothesized that the low-to-moderate PM localization efficiency of the original mgcBCP/K6-SNAP_f_ SLIPT system resulted from cellular cleavage of the myrGC motif in mgcBCP, as observed for mgcTMP. When we investigated the possible cellular degradation of mgcBCP by HPLC and mass spectroscopy, we detected a cleavage product of mgcBCP lacking the myrGC motif (**Fig. S1**). We therefore synthesized a new SL **1**, replacing the myrGC motif of mgcBCP with the protease-resistant m^D^c motif (m^D^cBCP) (**Fig. 1b**). For the SLIPT assay, we used HeLa cells expressing K6-SNAP_f_-Halo (**Fig. 2a**) labeled with the HaloTag^®^ TMR (tetramethylrhodamine) ligand^14^ [K6-SNAP_f_-Halo(TMR)]. We evaluated the PM recruitment efficiency by quantifying the ratio of the PM to cytosolic fluorescence intensity (P/C ratio) of the protein after protein translocation induction. Whereas the original mgcBCP resulted in a P/C ratio of 1.3 ± 0.1 (**Fig. 2b,f**), the new SL, m^D^cBCP, induced a significantly improved PM recruitment of K6-SNAP_f_-Halo(TMR) with a P/C ratio of 1.6 ± 0.1 (**Fig. 2c,f**).^15^

**Fig. 2.**
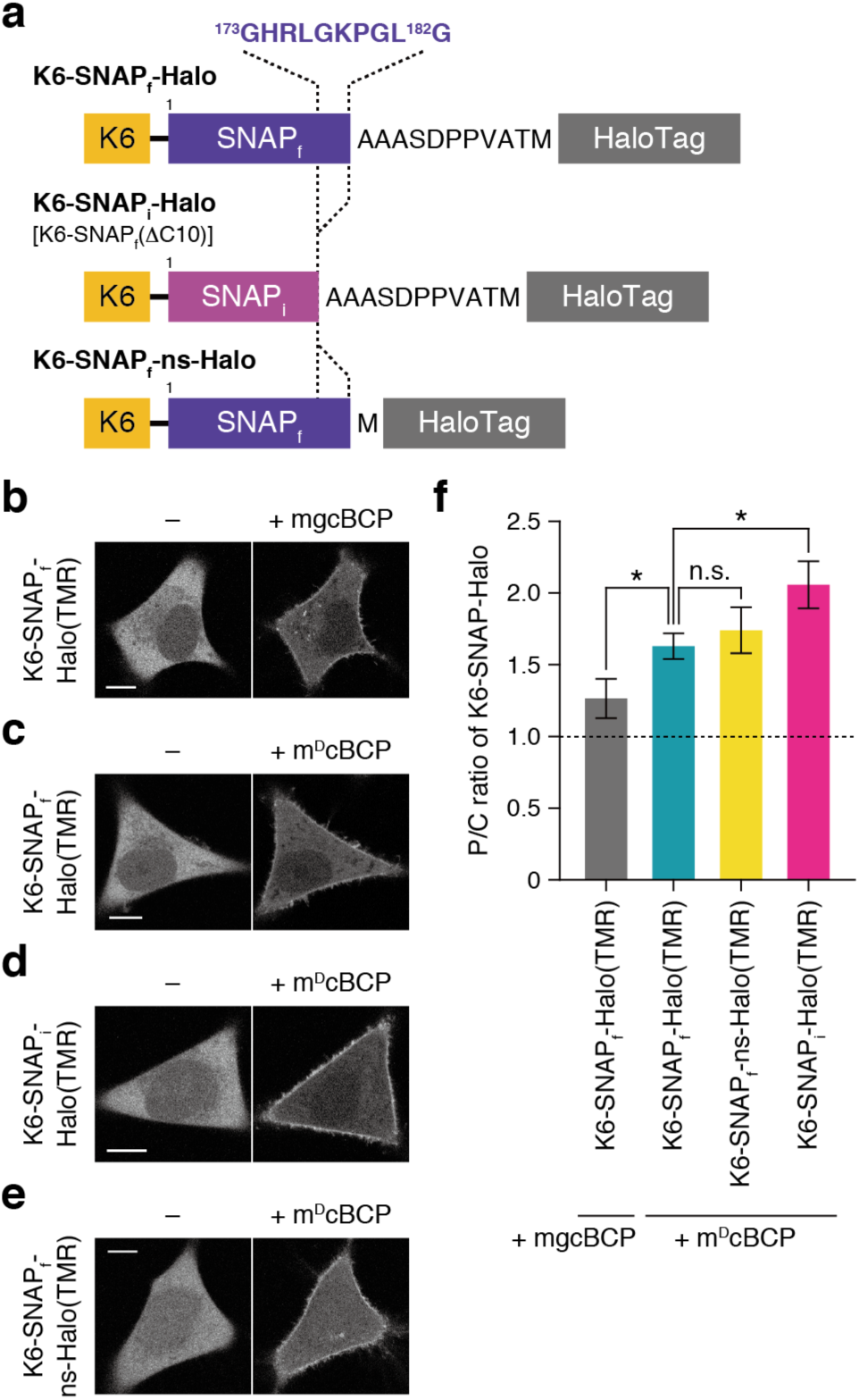
Development of an improved SNAP SLIPT system. (**a**) Schematic representations of the K6-tagged SNAP-HaloTag fusions (ns, no spacer). (**b**) Translocation of K6-SNAPf-Halo(TMR) by mgcBCP. (**c**–**e**) Translocation of K6-SNAPf-Halo(TMR) (**c**), K6-SNAP_f_(ΔC10)-Halo(TMR) (**d**), and K6-SNAPf-ns-Halo(TMR) (**e**) by m^D^cBCP (**1**). For **b**–**e**, confocal fluorescence images of HeLa cells expressing the indicated construct were taken before (left) and 60 min after incubation with the indicated SL (10 µM) (right). Scale bars, 10 µm. (**f**) Quantification of the PM translocation efficiency. The ratios of the PM to the cytosolic fluorescence intensity (P/C ratios) of the indicated construct were quantified after treatment with the indicated SL (10 µM) for 60 min. Cells with similar expression levels were used. Data are presented as the mean ± SD (n ≥ 9 cells). The symbols indicate the results of *t* test analysis; n.s.: *p* > 0.05, \**p* < 0.05.

We next attempted to improve the performance of the K6-SNAP_f_ SLIPT system by engineering the K6-SNAP_f_ domain. Three crystal structures of SNAP-tag (PDB ID: 3KZZ)^16^ and its ancestor human *O*^6^-alkylguanine-DNA alkyltransferase (PDB IDs: 1QNT^17^ and 1EH6^18^) together suggest that the C-terminal polypeptide chain following the H5 helix is oriented such that the C-terminus (the protein fusion site) points to the PM when the protein is anchored on the PM by SL-mediated lipidation (**Fig. S2**). If this is true, protein fusion to the SNAP_f_ C-terminus may cause steric hindrance and impede the binding of the lipidated K6-SNAP_f_-fusion protein to the PM. Based on this hypothesis, we attempted to construct and test a K6-SNAP_f_-Halo variant in which the C-terminal 10 amino acids of SNAP_f_ (^173^GHRLGKPGL^182^G) were deleted [K6-SNAP_f_(ΔC10)-Halo] (**Fig. 2a**). Interestingly, the K6-SNAP_f_(ΔC10)-Halo(TMR) showed further improvement in PM translocation by m^D^cBCP with a P/C ratio of 2.1 ± 0.2 (**Fig. 2d,f**). In contrast, when we deleted a 10-amino-acid linker of the original K6-SNAP_f_-Halo (K6-SNAP_f_-ns-Halo) (**Fig. 2a**) instead of the C-terminal 10 amino acids of SNAP_f_, no significant improvement of the PM translocation efficiency was observed (P/C ratio = 1.7 ± 0.2) (**Fig. 2e,f**). These results indicate that the enhanced PM translocation efficiency was caused by deleting the C-terminal 10 amino acids of SNAP_f_, rather than shortening the linker between SNAP_f_ and HaloTag. Hereinafter, we refer to the improved K6-SNAP_f_(ΔC10) tag as K6-SNAP_i_ and use the m^D^cBCP/K6-SNAP_i_ pair as the second-generation PM-targeted SNAP SLIPT system.^18^

We next applied the m^D^cBCP/K6-SNAP_i_ pair for cell signal control. We first focused on the activation of the endogenous small GTPase Rac1 by recruiting Tiam1, a guanine nucleotide exchange factor for Rac1, to the PM (**Fig. 3a**).^6,20^ Activation of Rac1 leads to actin reorganization and lamellipodia formation. In our initial attempt, we used the original mgcBCP/K6-SNAP_f_ pair. We generated a fusion protein of K6-SNAP_f_-EGFP and Tiam1 (K6S_f_G-Tiam1) and expressed this protein in HeLa cells. Owing to the low protein recruitment efficiency of the original system, no significant morphological change of the cells was observed after adding mgcBCP (**Fig. 3c**). In contrast, the same experiment using the new m^D^cBCP/K6-SNAP_i_ pair induced a significant formation of thin lamellipodia along the periphery of the HeLa cells expressing K6-SNAP_i_-EGFP-Tiam1 (K6S_i_G-Tiam1), indicating the efficient activation of Rac1 (**Fig. 3b,c**). No such lamellipodia formation occurred with the PM recruitment of K6-SNAP_i_-EGFP lacking Tiam1 (K6S_i_G) (**Fig. 3c**). In comparison with the original system, these results clearly demonstrate the superior performance of the m^D^cBCP/K6-SNAP_i_-based SLIPT system as a tool for synthetic cell signal manipulation. Furthermore, the new system could be used to control other signaling molecules, including cRaf (**Fig. S4**), PI3K (**Fig. S5**), and Sos (**Fig. S6**), thereby verifying the broad applicability of the system.

**Fig. 3.**
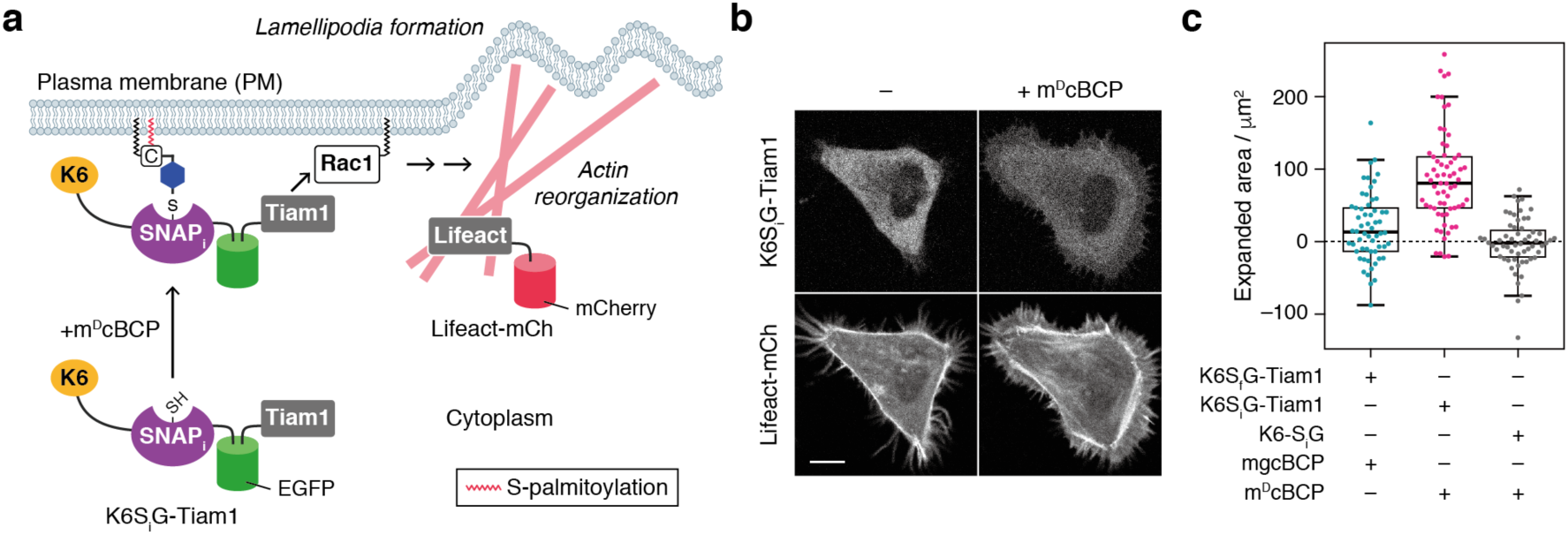
Synthetic Tiam1-mediated Rac1 activation and lamellipodia formation in HeLa cells by the second-generation SNAP SLIPT system. (**a**) Schematic illustration of the experimental setup for Tiam1-mediated Rac1 activation. (**b**) Confocal fluorescence images of HeLa cells coexpressing K6-SNAPi-EGFP-Tiam1 (K6SiG-Tiam1) and Lifeact-mCherry^21^ (Lifeact-mCh) were taken before (left) and 60 min after incubation with 10 μM m^D^cBCP (right). Scale bar, 10 µm. (**c**) Quantification of the expanded cell area. The Rac1 activation assay was performed using the SNAP-fusion/SL pairs shown in the graph. The expanded area was calculated by determining the cell area difference before chemical treatment and after incubation with the indicated SL (10 μM) for 60 min (n ≥ 58 cells). Central lines represent the median values.

In conclusion, by redesigning the SL and protein tag, we developed a versatile second-generation SNAP SLIPT system based on the m^D^cBCP/K6-SNAP_i_ pair. This system enables efficient and rapid PM-specific translocation of various K6-SNAP_i_-fusion proteins and cell signal manipulation in a time scale of minutes based on synthetic post-translational protein lipidation. Because various lipid and lipopeptide structures (including unnatural analogs) can be covalently attached to the target protein, this synthetic post-translational protein lipidation approach could also potentially be used to investigate how the lipid/lipopeptide structure regulates the localization and function of lipidated proteins in cells. In combination with the previously established eDHFR SLIPT system, the present SNAP SLIPT system may be a valuable research tool in chemical biology and synthetic biology for manipulating multiple signaling molecules in living single cells.^8^

## Supplementary information available

Supplementary figures S1–S8, synthesis and characterization of compounds, and supplementary methods for molecular and cell biology experiments.

## Conflicts of interest

T.Y., S.S., and S.T. are co-inventors on Japan patent application No. 2020-030868 that includes the m^D^c motif described in this paper. Other authors declare no competing interests.

## Acknowledgment

We thank Dr. Akinobu Nakamura (National Institute for Basic Biology) for technical assistance. This work was supported by JSPS Grants-in-Aid for Scientific Research (KAKENHI) (15H03835, 15H05949 “Resonance Bio”, 18H02086, and 18H04546 “Chemistry for Multimolecular Crowding Biosystems”), the Uehara Memorial Foundation, and the Takeda Science Foundation (to S.T.). This work was also supported in part by MEXT Leading Initiative for Excellent Young Researchers and by JST PRESTO (JPMJPR178B) (to T.Y.). S.S. acknowledges scholarship support from the Hirota Scholarship Society and the SUNBOR Scholarship from the Suntory Foundation for Life Sciences.

## Notes

### Competing Interest Statement

T.Y., S.S., and S.T. are co-inventors on Japan patent application No. 2020-030868 that includes the mDc motif described in this paper. Other authors declare no competing interests.

## Notes and references

1. M. D. Resh, Biochim. Biophys. Acta, 1999, 1451, 1–16.

2. M. D. Resh, Nat. Chem. Biol., 2006, 2, 584–590.

3. M. E. Linder and R. J. Deschenes, Nat. Rev. Mol. Cell Biol., 2007, 8, 74–84.

4. M. Wang and P. J. Casey, Nat. Rev. Mol. Cell Biol., 2016, 17, 110–122.

5. In a pioneering work, Rudd and coworkers demonstrated the spatiotemporally controlled anchoring of SNAP-tag to phospholipid membranes (liposomes) based on covalent lipid modification: A. K. Rudd, J. M. V. Cuevas and N. K. Devaraj, J. Am. Chem. Soc., 2015, 137, 4884–4887.

6. M. Ishida, H. Watanabe, K. Takigawa, Y. Kurishita, C. Oki, A. Nakamura, I. Hamachi and S. Tsukiji, J. Am. Chem. Soc., 2013, 135, 12684–12689.

7. A. Nakamura, R. Katahira, S. Sawada, E. Shinoda, K. Kuwata, T. Yoshii and S. Tsukiji, Biochemistry, 2020, 59, 205–211.

8. A. Nakamura, C. Oki, K. Kato, S. Fujinuma, G. Maryu, K. Kuwata, T. Yoshii, M. Matsuda, K. Aoki and S. Tsukiji, ACS Chem. Biol., Article ASAP, DOI: 10.1021/acschembio.0c00024.

9. A. Nakamura, C. Oki, S. Sawada, T. Yoshii, K. Kuwata, A. K. Rudd, N. K. Devaraj, K. Noma and S. Tsukiji, ACS Chem. Biol., Article ASAP, DOI: 10.1021/acschembio.0c00014.

10. A. Keppler, S. Gendreizig, T. Gronemeyer, H. Pick, H. Vogel and K. Jonsson, Nat. Biotechnol., 2003, 21, 86–89.

11. X. Sun, A. Zhang, B. Baker, L. Sun, A. Howard, J. Buswell, D. Maurel, A. Masharina, K. Johnsson, C. J. Noren, M.-Q. Xu and I. R. Corrêa, Jr., ChemBioChem, 2011, 12, 2217–2226.

12. S. P. Creaser and B. R. Peterson, J. Am. Chem. Soc., 2002, 124, 2444–2445.

13. H. Schroeder, R. Leventis, S. Shahinian, P. A. Walton and J. R. Silvius, J. Cell Biol., 1996, 134, 647–660.

14. G. V. Los, L. P. Encell, M. G. McDougall, D. D. Hartzell, N. Karassina, C. Zimprich, M. G. Wood, R. Learish, R. F. Ohana, M. Urh, D. Simpson, J. Mendez, K. Zimmerman, P. Otto, G. Vidugiris, J. Zhu, A. Darzins, D. H. Klaubert, R. F. Bulleit and K. V. Wood, ACS Chem. Biol., 2008, 3, 373–382.

15. In agreement with our hypothesis, no degradation product of m^D^cBCP was detected by HPLC (Fig. S1).

16. B. Mollwitz, E. Brunk, S. Schmitt, F. Pojer, M. Bannwarth, M. Schiltz, U. Rothlisberger and K. Johnsson, Biochemistry, 2012, 51, 986–994.

17. J. E. A. Wibley, A. E. Pegg and P. C. E. Moody, Nucleic Acids Res., 2000, 28, 393–401.

18. D. S. Daniels, C. D. Mol, A. S. Arvai, S. Kanugula, A. E. Pegg and J. A. Tainer, EMBO J., 2000, 19, 1719–1730.

19. The m^D^cBCP/K6-SNAP_i_pair also induced a more efficient PM translocation of an EGFP fusion compared with the original mgcBCP/K6-SNAP_f_system (Fig. S3).

20. T. Inoue, W. D. Heo, J. S. Grimley, T. J. Wandless and T. Meyer, Nat. Methods, 2005, 2, 415–418.

21. J. Riedl, A. H. Crevenna, K. Kessenbrock, J. H. Yu, D. Neukirchen, M. Bista, F. Bradke, D. Jenne, T. A. Holak, Z. Werb, M. Sixt and R. Wedlich-Soldner, Nat. Methods, 2008, 5, 605–607.

