## Supplementary Information for "Improved synthetic lipidation-based protein translocation system for SNAP-tag fusion proteins"

### Supplementary Figures

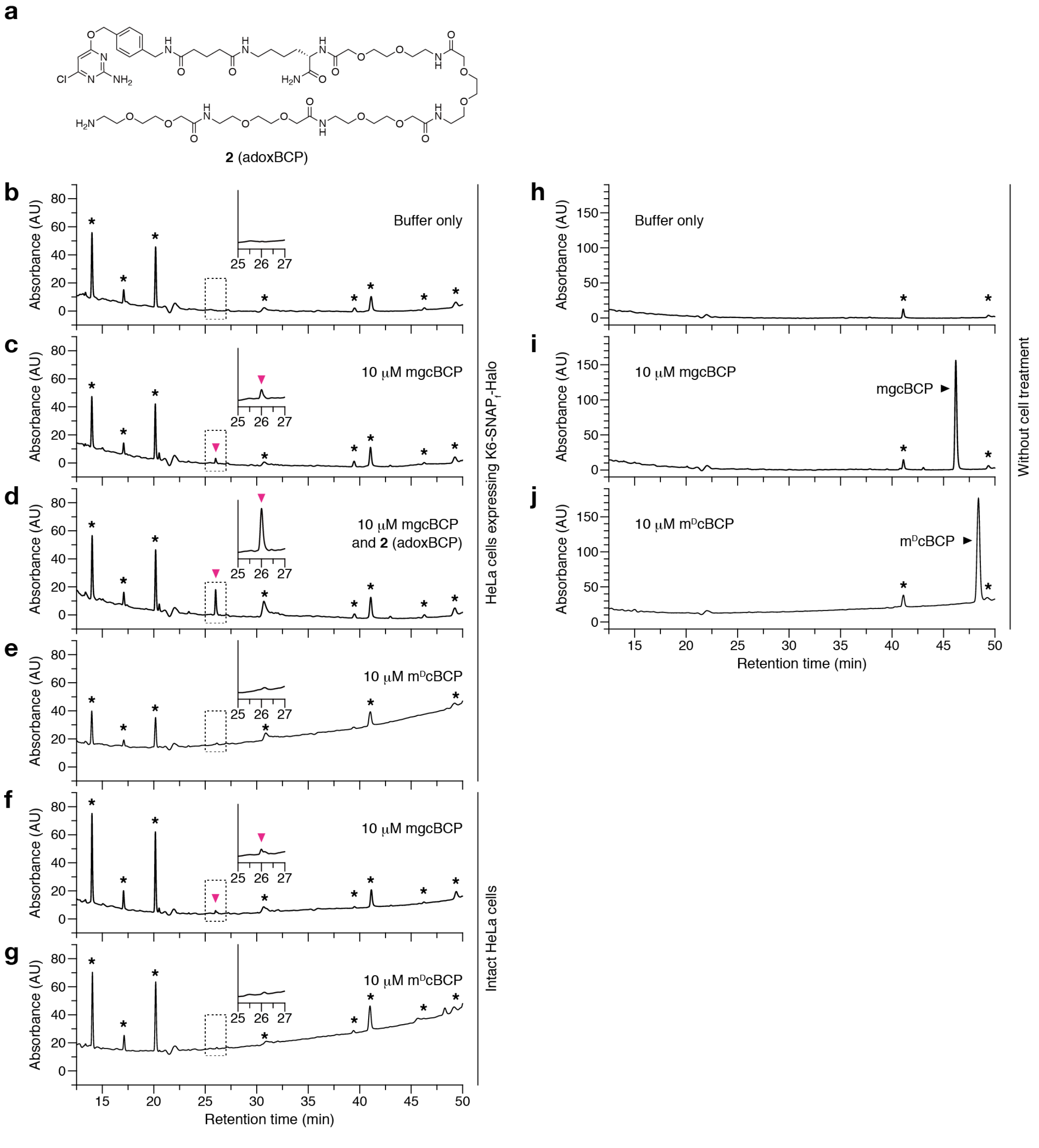


**Fig. S1.** HPLC analysis of mgcBCP cleavage. (**a**) Chemical structure of compound **2** (adoxBCP). (**b**) HPLC chart of HEPES buffered saline (HBS) (no ligand) collected after incubation with K6-SNAP$_f$-Halo-expressing HeLa cells at 37 °C for 3 h. (**c**) HPLC chart of mgcBCP (10 $\mu$M)-containing HBS collected after incubation with K6-SNAP$_f$-Halo-expressing HeLa cells at 37 °C for 3 h. (**d**) HPLC chart of a mixture of mgcBCP (10 $\mu$M)-containing HBS treated as described in (**c**) and chemically synthesized compound **2**. (**e**) HPLC chart of m$^D$cBCP (10 $\mu$M)-containing HBS treated as described in (**c**). (**f**) HPLC chart of mgcBCP (10 $\mu$M)-containing HBS collected after incubation with intact HeLa cells at 37 °C for 3 h. (**g**) HPLC chart of m$^{D}$cBCP (10 $\mu$M)-containing HBS treated as described in (**f**). (**h**) HPLC chart of HBS alone (no ligand and no cell treatment). (**i**) HPLC chart of mgcBCP (10 $\mu$M)-containing HBS incubated at 37 °C for 3 h (without cells). (**j**) HPLC chart of m$^{D}$cBCP (10 $\mu$M)-containing HBS incubated at 37 °C for 3 h (without cells). Asterisks indicate background peaks. In **c**, the new peak (around 26 min) indicated by a pink arrowhead was collected and analyzed by high-resolution mass spectrometry. The observed mass was consistent with the theoretical mass of compound **2** (adoxBCP): calculated for $[M+H]^+$, 1231.5973; found, 1231.5924. In **d**, the new peak overlapped with the chemically synthesized adoxBCP. These results demonstrate that the amide bond C-terminal to the Cys residue of mgcBCP was cleaved during the incubation with the cells. No such degradation was observed in the case of m$^{D}$cBCP (**e**,**g**).

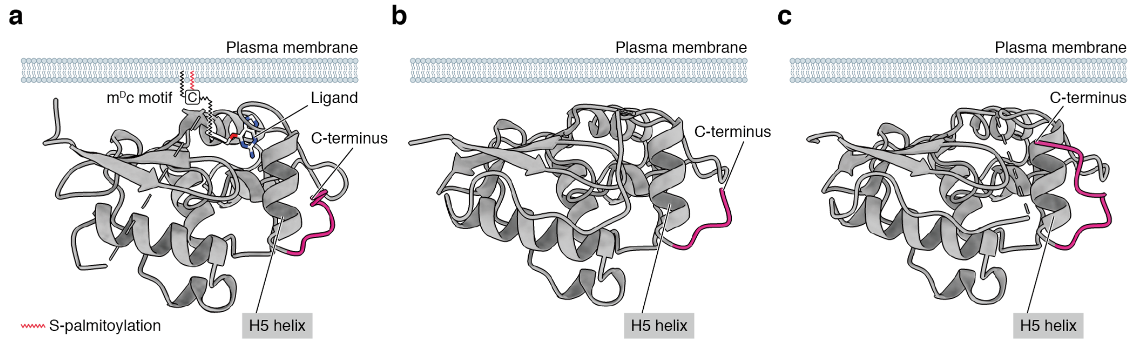


**Fig. S2.** The C-terminus of SNAP-tag points to the PM when it is anchored on the PM by $m^{D}$cBCP. (**a**) Crystal structure of SNAP-tag bound to its substrate benzylguanine (PDB: 3KZZ[S1]). The structure is depicted in a manner that the SNAP-tag is anchored on the putative PM by $m^{D}$cBCP. (**b**,**c**) Crystal structures of human $O^{6}$-alkylguanine-DNA alkyltransferase [PDB IDs: 1QNT[S2] (**b**) and 1EH6[S3] (**c**)]. The C-terminal polypeptide chain following the H5 helix is shown in magenta. All three crystal structures suggest that the C-terminus of SNAP-tag points to the PM in this membrane-anchored orientation. These models were displayed with UCSF ChimeraX.[S4]

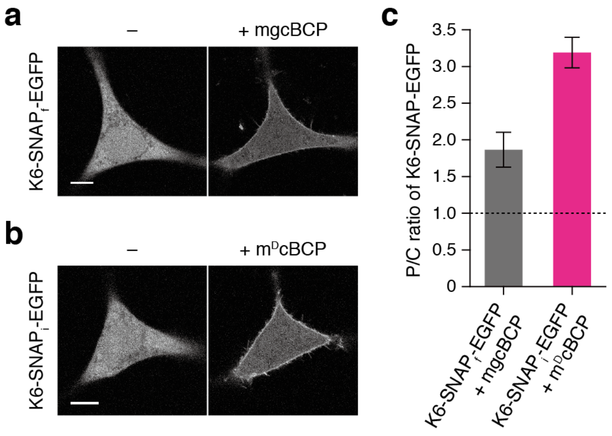


**Fig. S3** PM translocation of EGFP fusion proteins by the original and second-generation SLIPT systems. (**a**,**b**) Confocal fluorescence images of HeLa cells expressing K6-SNAP$_f$-EGFP (**a**) and K6-SNAP$_i$-EGFP (**b**) were taken before (left) and 60 min after incubation with the indicated SL (10 µM) (right). Scale bars, 10 µm. (**c**) Quantification of the PM translocation efficiency. The ratios of the PM to the cytosolic fluorescence intensity (P/C ratios) of the indicated construct were quantified after treatment with the indicated SL (10 µM) for 60 min. Cells with similar expression levels were used. Data are presented as the mean ± SD (n = 6 cells).

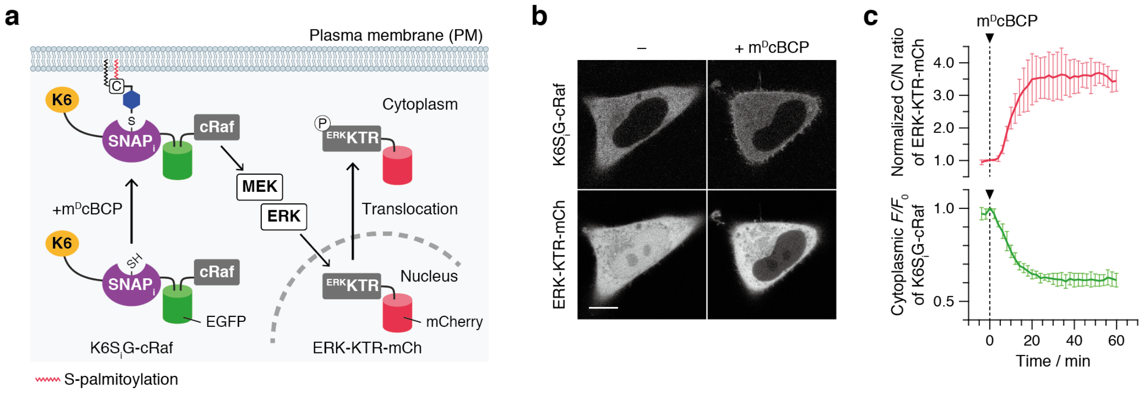


**Fig. S4.** Synthetic cRaf-mediated ERK activation by the second-generation SNAP SLIPT system. (**a**) Schematic illustration of the experimental setup for Raf-mediated ERK activation.[S5] (**b**) Confocal fluorescence images of HeLa cells coexpressing K6-SNAP$_i$-EGFP-cRaf (K6S$_i$G-cRaf) and ERK-KTR-mCherry[S6] (ERK-KTR-mCh) were taken before (left) and 60 min after incubation with 10 μM m$^D$cBCP (right). Scale bars, 10 μm. (**c**) Time course of K6S$_i$G-cRaf translocation and ERK activation. To evaluate the K6S$_i$G-cRaf translocation (green), normalized fluorescence intensities of K6S$_i$G-cRaf in the cytoplasm were plotted as a function of time. To evaluate the ERK activity (red), normalized ratios of the cytoplasm to the nuclear fluorescence intensity (C/N ratios) of ERK-KTR-mCh were plotted as a function of time. Data are presented as the mean ± SD (n = 4 cells).

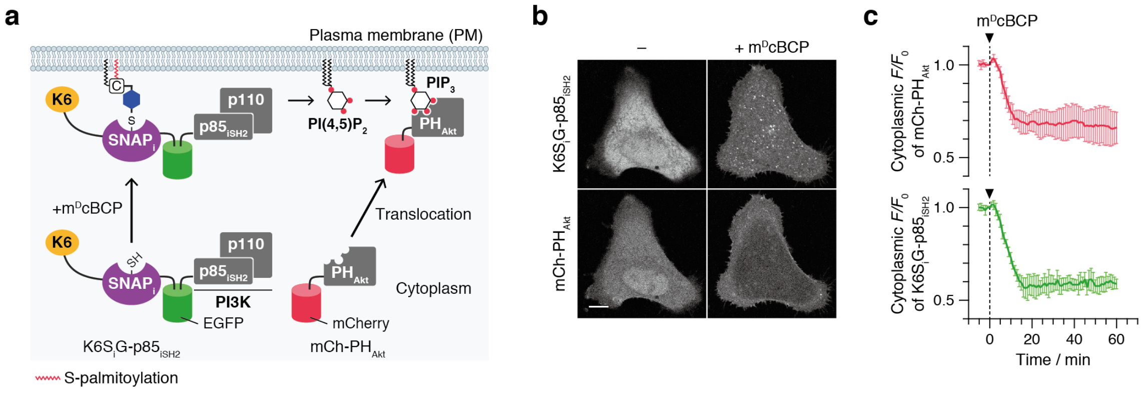


**Fig. S5.** Synthetic PI3K-mediated PI(3,4,5)$P_3$ production by the second-generation SNAP SLIPT system. (**a**) Schematic illustration of the experimental setup for the PI3K-mediated PI(3,4,5)$P_3$ production.[S7] (**b**) Confocal fluorescence images of HeLa cells coexpressing K6-SNAP$_i$-EGFP-p85$_{iSH2}$ (K6S$_i$G-p85$_{iSH2}$) and mCherry-PH$_{Akt}$[S8] (mCh-PH$_{Akt}$) were taken before (left) and 60 min after incubation with 10 µM m$^D$cBCP (right). Scale bars, 10 $\mu$m. (**c**) Time course of K6S$_i$G-p85$_{iSH2}$ translocation and PI(3,4,5)$P_3$ production. To evaluate the K6S$_i$G-p85$_{iSH2}$ translocation (green), normalized fluorescence intensities of K6S$_i$G-p85$_{iSH2}$ in the cytoplasm were plotted as a function of time. To evaluate the PI(3,4,5)$P_3$ production (red), normalized fluorescence intensities of mCh-PH$_{Akt}$ in the cytoplasm were plotted as a function of time. Data are presented as the mean ± SD (n = 4 cells).

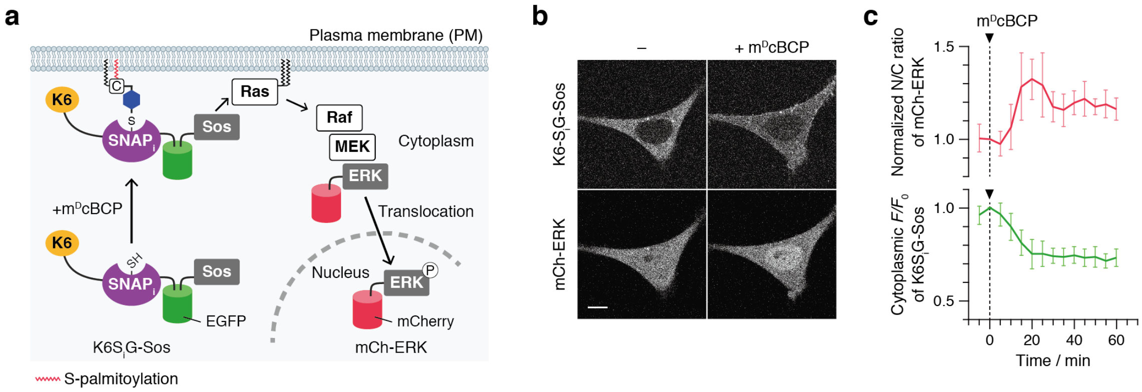


**Fig. S6.** Synthetic Sos-mediated Ras/ERK activation by the second-generation SNAP SLIPT system. (**a**) Schematic illustration of the experimental setup for the Sos-mediated Ras/ERK activation.[S9] (**b**) Confocal fluorescence images of HeLa cells coexpressing K6-SNAP$_i$-EGFP-Sos (K6S$_i$G-Sos) and mCherry-ERK[S10] (mCh-ERK) were taken before (left) and 20 min after incubation with 10 µM m$^D$cBCP (right). Scale bars, 10 $\mu$m. (**c**) Time course of K6S$_i$G-Sos translocation and ERK activation. To evaluate the K6S$_i$G-Sos translocation (green), normalized fluorescence intensities of K6S$_i$G-Sos in the cytoplasm were plotted as a function of time. To evaluate the ERK activity (red), normalized ratios of the nuclear to cytoplasm fluorescence intensity (N/C ratios) of mCh-ERK were plotted as a function of time. Data are presented as the mean ± SD (n = 6 cells).

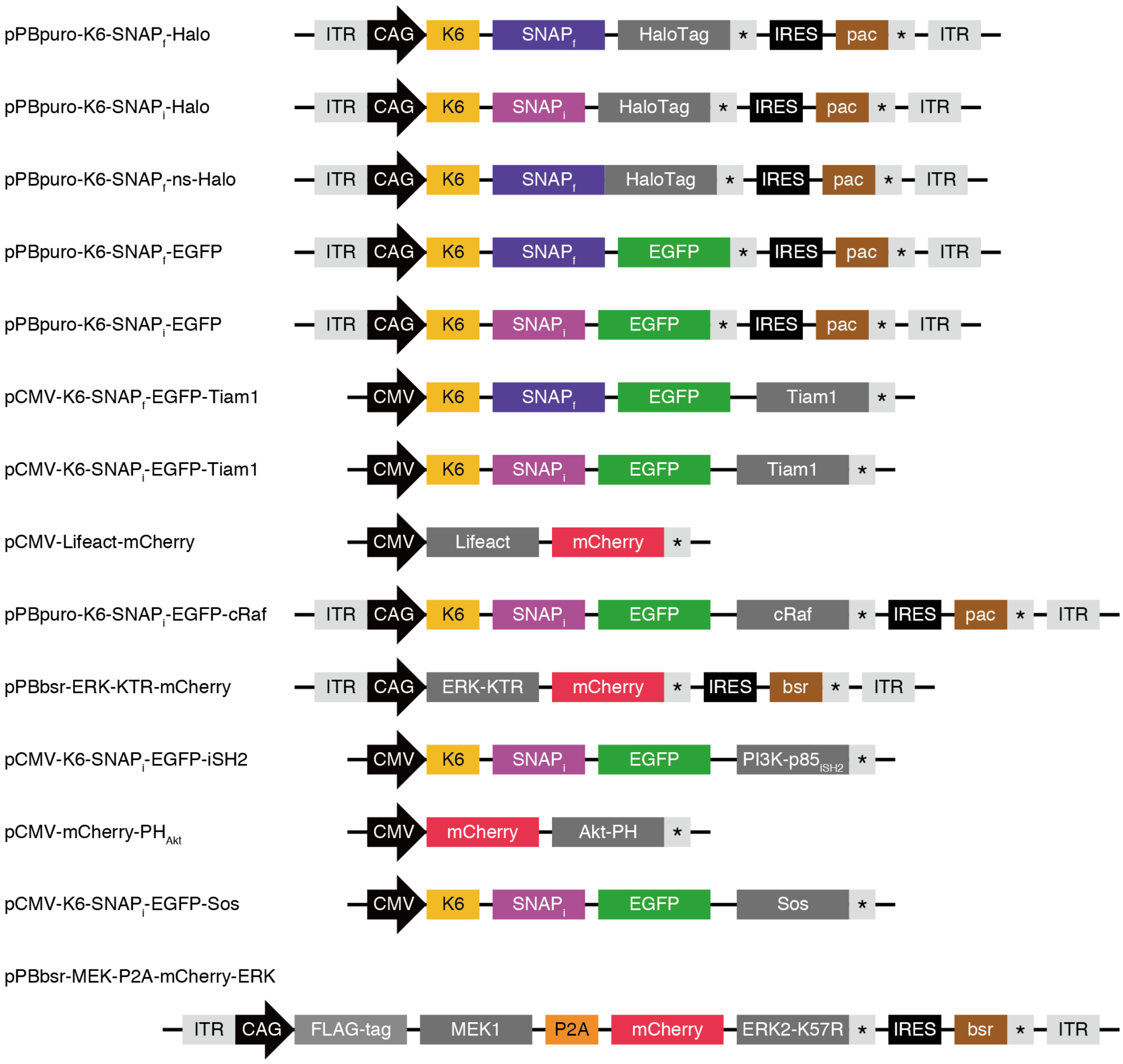


**Fig. S7** Schematic illustration of the domain structures of constructs used in this study.

**a**

pPBpuro-K6-SNAP$_f$-Halo

```
>Amino acid sequence
MKKKKKKGSGADKDCEMKRTTLDSPLGKLELSGCEQGLHRIIFLGKGTSAADAVEVPAPAA
VLGGPEPLMQATAWLNAYFHQPEAIEEFPVPALHHPVFQQESFTRQVLWKLLKVVKFGEVI
SYSHLAALAGNPAATAAVKTALSGNPVPILIPCHRVVQGDLDVGGYEGGLAVKEWLLAHEG
HRLGKPGLGAAASDPPVATMAEIGTGFPFDPHYVEVLGERMHYVDVGPRDGTPVLFLHGNP
TSSYVWRNIIPHVAPTHRCIAPDLIGMGKSDKPDLGYFFDDHVRFMDAFIEALGLEEVVLV
IHDWGSALGFHWAKRNPERVKGIAFMEFIRPIPTWDEWPEFARETFQAFRTTDVGRKLIID
QNVFIEGTLPMGVVRPLTEVEMDHYREPFLNPVDREPLWRFPNELPIAGEPANIVALVEEY
MDWLHQSPVPKLLFWGTPGVLIPPAEAARLAKSLPNCKAVDIGPGLNLLQEDNPDLIGSEI
ARWLSTLEISGLYK*-[EMCV  IRES]-MTEYKPTVRLATRDDVPRAVRTLAAAFADYPAT
RHTVDPDRHIERVTELQELFLTRVGLDIGKVWVADDGAAVAVWTTPESVEAGAVFAEIGPR
MAELSGSRLAAQQQMEGLLAPHRPKEPAWFLATVGVSPDHQGKGLGSAVVLPGVEAAERAG
VPAFLETSAPRNLPFYERLGFTVTADVECPKDRATWCMTRKPGA*

>DNA sequence
ATGAAAAAAAAGAAAAAGAAAGGCAGCGGTGCTGACAAAGACTGCGAAATGAAGCGCACCA
CCCTGGATAGCCCTCTGGGCAAGCTGGAACTGTCTGGGTGCGAACAGGGCCTGCACCGTAT
CATCTTCCTGGGCAAAGGAACATCTGCCGCCGACGCCGTGGAAGTGCCTGCCCCAGCCGCC
GTGCTGGGCGGACCAGAGCCACTGATGCAGGCCACCGCCTGGCTCAACGCCTACTTTCACC
AGCCTGAGGCCATCGAGGAGTTCCCTGTGCCAGCCCTGCACCACCCAGTGTTCCAGCAGGA
GAGCTTTACCCGCCAGGTGCTGTGGAAACTGCTGAAAGTGGTGAAGTTCGGAGAGGTCATC
AGCTACAGCCACCTGGCCGCCCTGGCCGGCAATCCCGCCGCCACCGCCGCCGTGAAAACCG
CCCTGAGCGGAAATCCCGTGCCCATTCTGATCCCCTGCCACCGGGTGGTGCAGGGCGACCT
GGACGTGGGGGGCTACGAGGGCGGGCTCGCCGTGAAAGAGTGGCTGCTGGCCCACGAGGGC
CACAGACTGGGCAAGCCTGGGCTGGGTGCGGCCGCTTCGGACCCACCGGTCGCCACCATGG
CAGAAATCGGTACTGGCTTTCCATTCGACCCCCATTATGTGGAAGTCCTGGGCGAGCGCAT
GCACTACGTCGATGTTGGTCCGCGCGATGGCACCCCTGTGCTGTTCCTGCACGGTAACCCG
ACCTCCTCCTACGTGTGGCGCAACATCATCCCGCATGTTGCACCGACCCATCGCTGCATTG
CTCCAGACCTGATCGGTATGGGCAAATCCGACAAACCAGACCTGGGTTATTTCTTCGACGA
CCACGTCCGCTTCATGGATGCCTTCATCGAAGCCCTGGGTCTGGAAGAGGTCGTCCTGGTC
ATTCACGACTGGGGCTCCGCTCTGGGTTTCCACTGGGCCAAGCGCAATCCAGAGCGCGTCA
AAGGTATTGCATTTATGGAGTTCATCCGCCCTATCCCGACCTGGGACGAATGGCCAGAATT
TGCCCGCGAGACCTTCCAGGCCTTCCGCACCACCGACGTCGGCCGCAAGCTGATCATCGAT
CAGAACGTTTTTATCGAGGGTACGCTGCCGATGGGTGTCGTCCGCCCGCTGACTGAAGTCG
AGATGGACCATTACCGCGAGCCGTTCCTGAATCCTGTTGACCGCGAGCCACTGTGGCGCTT
CCCAAACGAGCTGCCAATCGCCGGTGAGCCAGCGAACATCGTCGCGCTGGTCGAAGAATAC
ATGGACTGGCTGCACCAGTCCCCTGTCCCGAAGCTGCTGTTCTGGGGCACCCCAGGCGTTC
TGATCCCACCGGCCGAAGCCGCTCGCCTGGCCAAAAGCCTGCCTAACTGCAAGGCTGTGGA
CATCGGCCCAGGTCTGAATCTGCTGCAAGAAGACAACCCGGACCTGATCGGCAGCGAGATC
GCGCGCTGGCTGTCCACGCTGGAGATTTCCGGCCTGTACAAGTAAAGCGGCCGCGACTCTA
GATCATAATCAGCCATACCACATTTGTAGAGGTTTTACTTGCTTTAAAAAACCTCCCACAC
CTCCCCCTGAACCTGAAACATAAAATGAATGCAATTCTGCAGTCGACGGTACCGCGGGCCC
GGGATAAGTCAACTAACTTAAGCTAGCAACGGTTTCCCTCTAGCGGGATCAATTCCGcccc
cccccctaacgttactggccgaagccgcttggaataaggccggtgtgcgtttgtctatat
gttattttccaccatattgccgtcttttggcaatgtgagggcccggaaacctggccctgtc
ttcttgacgagcattcctaggggtctttcccctctcgccaaaggaatgcaaggtctgttga
atgtcgtgaaggaagcagttcctctggaagcttcttgaagacaaacaacgtctgtagcgac
cctttgcaggcagcggaaccccccacctggcgacaggtgcctctgcggccaaaagccacgt
gtataagatacacctgcaaaggcggcacaaccccagtgccacgttgtgagttggatagttg
tggaaagagtcaaatggctctcctcaagcgtattcaacaaggggctgaaggatgcccagaa
```

ggtaccccattgtatgggatctgatctggggcctcggtgcacatgctttacatgtgtttag
tcgaggttaaaaaacgtctaggccccccgaaccacggggacgtggttttcctttgaaaaac
acgatAATACCATGACCGAGTACAAGCCCACGGTGCGCCTCGCCACCCGCGACGACGTCCC
CAGGGCCGTACGCACCCTCGCCGCCGCGTTCGCCGACTACCCCGCCACGCGCCACACCGTC
GATCCGGACCGCCACATCGAGCGGGTCACCGAGCTGCAAGAACTCTTCCTCACGCGCGTCG
GGCTCGACATCGGCAAGGTGTGGGTCGCGGACGACGGCGCCGCGGTGGCGGTCTGGACCAC
GCCGGAGAGCGTCGAAGCGGGGGCGGTGTTCGCCGAGATCGGCCCGCGCATGGCCGAGTTG
AGCGGTTCCCGGCTGGCCGCGCAGCAACAGATGGAAGGCCTCCTGGCGCCGCACCGGCCCA
AGGAGCCCGCGTGGTTCCTGGCCACCGTCGGCGTCTCGCCCGACCACCAGGGCAAGGGTCT
GGGCAGCGCCGTCGTGCTCCCCGGAGTGGAGGCGGCCGAGCGCGCCGGGGTGCCCGCCTTC
CTGGAGACCTCCGCGCCCCGCAACCTCCCCTTCTACGAGCGGCTCGGCTTCACCGTCACCG
CCGACGTCGAGTGCCCGAAGGACCGCGCGACCTGGTGCATGACCCGCAAGCCCGGTGCCTG
A

**b**

pPBpuro-K6-SNAP$_i$-Halo

```
>Amino acid sequence
MKKKKKKGSGADKDCEMKRTTLDSPLGKLELSGCEQGLHRIIFLGKGTSAADAVEVPAPAA
VLGGPEPLMQATAWLNAYFHQPEAIEEFPVPALHHPVFQQESFTRQVLWKLLKVVKFGEVI
SYSHLAALAGNPAATAAVKTALSGNPVPILIPCHRVVQGDLDVGGYEGGLAVKEWLLAHEA
AASDPPVATMAEIGTGFPFDPHYVEVLGERMHYVDVGPRDGTPVLFLHGNPTSSYVWRNII
PHVAPTHRCIAPDLIGMGKSDKPDLGYFFDDHVRFMDAFIEALGLEEVVLVIHDWGSALGF
HWAKRNPERVKGIAFMEFIRPIPTWDEWPEFARETFQAFRTTDVGRKLIIDQNVFIEGTLP
MGVVRPLTEVEMDHYREPFLNPVDREPLWRFPNELPIAGEPANIVALVEEYMDWLHQSPVP
KLLFWGTPGVLIPPAEAARLAKSLPNCKAVDIGPGLNLLQEDNPDLIGSEIARWLSTLEIS
GLYK*-[EMCV IRES]-MTEYKPTVRLATRDDVPRAVRTLAAAFADYPATRHTVDPDRHI
ERVTELQELFLTRVGLDIGKVWVADDGAAVAVWTTPESVEAGAVFAEIGPRMAELSGSRLA
AQQQMEGLLAPHRPKEPAWFLATVGVSPDHQGKGLGSAVVLPGVEAAERAGVPAFLETSAP
RNLPFYERLGFTVTADVECPKDRATWCMTRKPGA*

>DNA sequence
ATGAAAAAAAAGAAAAAGAAAGGCAGCGGTGCTGACAAAGACTGCGAAATGAAGCGCACCA
CCCTGGATAGCCCTCTGGGCAAGCTGGAACTGTCTGGGTGCGAACAGGGCCTGCACCGTAT
CATCTTCCTGGGCAAAGGAACATCTGCCGCCGACGCCGTGGAAGTGCCTGCCCCAGCCGCC
GTGCTGGGCGGACCAGAGCCACTGATGCAGGCCACCGCCTGGCTCAACGCCTACTTTCACC
AGCCTGAGGCCATCGAGGAGTTCCCTGTGCCAGCCCTGCACCACCCAGTGTTCCAGCAGGA
GAGCTTTACCCGCCAGGTGCTGTGGAAACTGCTGAAAGTGGTGAAGTTCGGAGAGGTCATC
AGCTACAGCCACCTGGCCGCCCTGGCCGGCAATCCCGCCGCCACCGCCGCCGTGAAAACCG
CCCTGAGCGGAAATCCCGTGCCCATTCTGATCCCCTGCCACCGGGTGGTGCAGGGCGACCT
GGACGTGGGGGGCTACGAGGGCGGGCTCGCCGTGAAAGAGTGGCTGCTGGCCCACGAGGCG
GCCGCTTCGGACCCACCGGTCGCCACCATGGCAGAAATCGGTACTGGCTTTCCATTCGACC
CCCATTATGTGGAAGTCCTGGGCGAGCGCATGCACTACGTCGATGTTGGTCCGCGCGATGG
CACCCCTGTGCTGTTCCTGCACGGTAACCCGACCTCCTCCTACGTGTGGCGCAACATCATC
CCGCATGTTGCACCGACCCATCGCTGCATTGCTCCAGACCTGATCGGTATGGGCAAATCCG
ACAAACCAGACCTGGGTTATTTCTTCGACGACCACGTCCGCTTCATGGATGCCTTCATCGA
AGCCCTGGGTCTGGAAGAGGTCGTCCTGGTCATTCACGACTGGGGCTCCGCTCTGGGTTTC
CACTGGGCCAAGCGCAATCCAGAGCGCGTCAAAGGTATTGCATTTATGGAGTTCATCCGCC
CTATCCCGACCTGGGACGAATGGCCAGAATTTGCCCGCGAGACCTTCCAGGCCTTCCGCAC
CACCGACGTCGGCCGCAAGCTGATCATCGATCAGAACGTTTTTATCGAGGGTACGCTGCCG
ATGGGTGTCGTCCGCCCGCTGACTGAAGTCGAGATGGACCATTACCGCGAGCCGTTCCTGA
ATCCTGTTGACCGCGAGCCACTGTGGCGCTTCCCAAACGAGCTGCCAATCGCCGGTGAGCC
AGCGAACATCGTCGCGCTGGTCGAAGAATACATGGACTGGCTGCACCAGTCCCCTGTCCCG
AAGCTGCTGTTCTGGGGCACCCCAGGCGTTCTGATCCCACCGGCCGAAGCCGCTCGCCTGG
CCAAAAGCCTGCCTAACTGCAAGGCTGTGGACATCGGCCCAGGTCTGAATCTGCTGCAAGA
AGACAACCCGGACCTGATCGGCAGCGAGATCGCGCGCTGGCTGTCCACGCTGGAGATTTCC
GGCCTGTACAAGTAAAGCGGCCGCGACTCTAGATCATAATCAGCCATACCACATTTGTAGA
GGTTTTACTTGCTTTAAAAAACCTCCCACACCTCCCCCTGAACCTGAAACATAAAATGAAT
GCAATTCTGCAGTCGACGGTACCGCGGGCCCGGGATAAGTCAACTAACTTAAGCTAGCAAC
GGTTTCCCTCTAGCGGATCAATTCCGcccccccccccctaacgttactggccgaagccgct
tggaataaggccggtgtgcgtttgtctatatgttattttccaccatattgccgtcttttgg
caatgtgagggcccggaaacctggccctgtcttcttgacgagcattcctaggggtctttcc
cctctcgccaaaggaatgcaaggtctgttgaatgtcgtgaaggaagcagttcctctggaag
cttcttgaagacaaacaacgtctgtagcgaccctttgcaggcagcggaaccccccacctgg
cgacaggtgcctctgcggccaaaagccacgtgtataagatacacctgcaaaggcggcacaa
ccccagtgccacgttgtgagttggatagttgtggaaagagtcaatggctctcctcaagcg
tattcaacaaggggctgaaggatgcccagaaggtaccccattgtatgggatctgatctggg
```

gcctcggtgcacatgctttacatgtgtttagtcgaggttaaaaaacgtctaggccccccga
accacggggacgtggttttcctttgaaaaacacgatAATACCATGACCGAGTACAAGCCCA
CGGTGCGCCTCGCCACCCGCGACGACGTCCCCAGGGCCGTACGCACCCTCGCCGCCGCGTT
CGCCGACTACCCCGCCACGCGCCACACCGTCGATCCGGACCGCCACATCGAGCGGGTCACC
GAGCTGCAAGAACTCTTCCTCACGCGCGTCGGGCTCGACATCGGCAAGGTGTGGGTCGCGG
ACGACGGCGCCGCGGTGGCGGTCTGGACCACGCCGGAGAGCGTCGAAGCGGGGGCGGTGTT
CGCCGAGATCGGCCCGCGCATGGCCGAGTTGAGCGGTTCCCGGCTGGCCGCGCAGCAACAG
ATGGAAGGCCTCCTGGCGCCGCACCGGCCCAAGGAGCCCGCGTGGTTCCTGGCCACCGTCG
GCGTCTCGCCCGACCACCAGGGCAAGGGTCTGGGCAGCGCCGTCGTGCTCCCCGGAGTGGA
GGCGGCCGAGCGCGCCGGGGTGCCCGCCTTCCTGGAGACCTCCGCGCCCCGCAACCTCCCC
TTCTACGAGCGGCTCGGCTTCACCGTCACCGCCGACGTCGAGTGCCCGAAGGACCGCGCGA
CCTGGTGCATGACCCGCAAGCCCGGTGCCTGA

**c**

pPBpuro-K6-SNAP$_f$-ns-Halo

```
>Amino acid sequence
MKKKKKKGSGADKDCEMKRTTLDSPLGKLELSGCEQGLHRIIFLGKGTSAADAVEVPAPAA
VLGGPEPLMQATAWLNAYFHQPEAIEEFPVPALHHPVFQQESFTRQVLWKLLKVVKFGEVI
SYSHLAALAGNPAATAAVKTALSGNPVPILIPCHRVVQGDLDVGGYEGGLAVKEWLLAHEG
HRLGKPGLGMAEIGTGFPFDPHYVEVLGERMHYVDVGPRDGTPVLFLHGNPTSSYVWRNII
PHVAPTHRCIAPDLIGMGKSDKPDLGYFFDDHVRFMDAFIEALGLEEVVLVIHDWGSALGF
HWAKRNPERVKGIAFMEFIRPIPTWDEWPEFARETFQAFRTTDVGRKLIIDQNVFIEGTLP
MGVVRPLTEVEMDHYREPFLNPVDREPLWRFPNELPIAGEPANIVALVEEYMDWLHQSPVP
KLLFWGTPGVLIPPAEAARLAKSLPNCKAVDIGPGLNLLQEDNPDLIGSEIARWLSTLEIS
GLYK*-[EMCV  IRES]-MTEYKPTVRLATRDDVPRAVRTLAAAFADYPATRHTVDPDRHI
ERVTELQELFLTRVGLDIGKVWVADDGAAVAVWTTPESVEAGAVFAEIGPRMAELSGSRLA
AQQQMEGLLAPHRPKEPAWFLATVGVSPDHQGKGLGSAVVLPGVEAAERAGVPAFLETSAP
RNLPFYERLGFTVTADVECPKDRATWCMTRKPGA*

>DNA sequence
ATGAAAAAAAAGAAAAAGAAAGGCAGCGGTGCTGACAAAGACTGCGAAATGAAGCGCACCA
CCCTGGATAGCCCTCTGGGCAAGCTGGAACTGTCTGGGTGCGAACAGGGCCTGCACCGTAT
CATCTTCCTGGGCAAAGGAACATCTGCCGCCGACGCCGTGGAAGTGCCTGCCCCAGCCGCC
GTGCTGGGCGGACCAGAGCCACTGATGCAGGCCACCGCCTGGCTCAACGCCTACTTTCACC
AGCCTGAGGCCATCGAGGAGTTCCCTGTGCCAGCCCTGCACCACCCAGTGTTCCAGCAGGA
GAGCTTTACCCGCCAGGTGCTGTGGAAACTGCTGAAAGTGGTGAAGTTCGGAGAGGTCATC
AGCTACAGCCACCTGGCCGCCCTGGCCGGCAATCCCGCCGCCACCGCCGCCGTGAAAACCG
CCCTGAGCGGAAATCCCGTGCCCATTCTGATCCCCTGCCACCGGGTGGTGCAGGGCGACCT
GGACGTGGGGGGCTACGAGGGCGGGCTCGCCGTGAAAGAGTGGCTGCTGGCCCACGAGGGC
CACAGACTGGGCAAGCCTGGGCTGGGTATGGCAGAAATCGGTACTGGCTTTCCATTCGACC
CCCATTATGTGGAAGTCCTGGGCGAGCGCATGCACTACGTCGATGTTGGTCCGCGCGATGG
CACCCCTGTGCTGTTCCTGCACGGTAACCCGACCTCCTCCTACGTGTGGCGCAACATCATC
CCGCATGTTGCACCGACCCATCGCTGCATTGCTCCAGACCTGATCGGTATGGGCAAATCCG
ACAAACCAGACCTGGGTTATTTCTTCGACGACCACGTCCGCTTCATGGATGCCTTCATCGA
AGCCCTGGGTCTGGAAGAGGTCGTCCTGGTCATTCACGACTGGGGCTCCGCTCTGGGTTTC
CACTGGGCCAAGCGCAATCCAGAGCGCGTCAAAGGTATTGCATTTATGGAGTTCATCCGCC
CTATCCCGACCTGGGACGAATGGCCAGAATTTGCCCGCGAGACCTTCCAGGCCTTCCGCAC
CACCGACGTCGGCCGCAAGCTGATCATCGATCAGAACGTTTTTATCGAGGGTACGCTGCCG
ATGGGTGTCGTCCGCCCGCTGACTGAAGTCGAGATGGACCATTACCGCGAGCCGTTCCTGA
ATCCTGTTGACCGCGAGCCACTGTGGCGCTTCCCAAACGAGCTGCCAATCGCCGGTGAGCC
AGCGAACATCGTCGCGCTGGTCGAAGAATACATGGACTGGCTGCACCAGTCCCCTGTCCCG
AAGCTGCTGTTCTGGGGCACCCCAGGCGTTCTGATCCCACCGGCCGAAGCCGCTCGCCTGG
CCAAAAGCCTGCCTAACTGCAAGGCTGTGGACATCGGCCCAGGTCTGAATCTGCTGCAAGA
AGACAACCCGGACCTGATCGGCAGCGAGATCGCGCGCTGGCTGTCCACGCTGGAGATTTCC
GGCCTGTACAAGTAAAGCGGCCGCGACTCTAGATCATAATCAGCCATACCACATTTGTAGA
GGTTTTACTTGCTTTAAAAAACCTCCCACACCTCCCCCTGAACCTGAAACATAAAATGAAT
GCAATTCTGCAGTCGACGGTACCGCGGGCCCGGGATAAGTCAACTAACTTAAGCTAGCAAC
GGTTTCCCTCTAGCGGGATCAATTCCGcccccccccctaacgttactggccgaagccgct
tggaataaggccggtgtgcgtttgtctatatgttattttccaccatattgccgtcttttgg
caatgtgagggcccggaaacctggccctgtcttcttgacgagcattcctaggggtctttcc
cctctcgccaaaggaatgcaaggtctgttgaatgtcgtgaaggaagcagttcctctggaag
cttcttgaagacaaacaacgtctgtagcgaccctttgcaggcagcggaaccccccacctgg
cgacaggtgcctctgcggccaaaagccacgtgtataagatacacctgcaaaggcggcacaa
ccccagtgccacgttgtgagttggatagttgtggaaagagtcaaatggctctcctcaagcg
tattcaacaaggggctgaaggatgcccagaaggtaccccattgtatgggatctgatctggg
```

gcctcggtgcacatgctttacatgtgtttagtcgaggttaaaaaacgtctaggccccccga
accacggggacgtggttttcctttgaaaaacacgatAATACCATGACCGAGTACAAGCCCA
CGGTGCGCCTCGCCACCCGCGACGACGTCCCCAGGGCCGTACGCACCCTCGCCGCCGCGTT
CGCCGACTACCCCGCCACGCGCCACACCGTCGATCCGGACCGCCACATCGAGCGGGTCACC
GAGCTGCAAGAACTCTTCCTCACGCGCGTCGGGCTCGACATCGGCAAGGTGTGGGTCGCGG
ACGACGGCGCCGCGGTGGCGGTCTGGACCACGCCGGAGAGCGTCGAAGCGGGGGCGGTGTT
CGCCGAGATCGGCCCGCGCATGGCCGAGTTGAGCGGTTCCCGGCTGGCCGCGCAGCAACAG
ATGGAAGGCCTCCTGGCGCCGCACCGGCCCAAGGAGCCCGCGTGGTTCCTGGCCACCGTCG
GCGTCTCGCCCGACCACCAGGGCAAGGGTCTGGGCAGCGCCGTCGTGCTCCCCGGAGTGGA
GGCGGCCGAGCGCGCCGGGGTGCCCGCCTTCCTGGAGACCTCCGCGCCCCGCAACCTCCCC
TTCTACGAGCGGCTCGGCTTCACCGTCACCGCCGACGTCGAGTGCCCGAAGGACCGCGCGA
CCTGGTGCATGACCCGCAAGCCCGGTGCCTGA

**d**

pPBpuro-K6-SNAP$_f$-EGFP

```
>Amino acid sequence
MKKKKKKGSGADKDCEMKRTTLDSPLGKLELSGCEQGLHRIIFLGKGTSAADAVEVPAPAA
VLGGPEPLMQATAWLNAYFHQPEAIEEFPVPALHHPVFQQESFTRQVLWKLLKVVKFGEVI
SYSHLAALAGNPAATAAVKTALSGNPVPILIPCHRVVQGDLDVGGYEGGLAVKEWLLAHEG
HRLGKPGLGAAASDPPVATMVSKGEELFTGVVPILVELDGDVNGHKFSVSGEGEGDATYGK
LTLKFICTTGKLPVPWPTLVTTLTYGVQCFSRYPDHMKQHDFFKSAMPEGYVQERTIFFKD
DGNYKTRAEVKFEGDTLVNRIELKGIDFKEDGNILGHKLEYNYNSHNVYIMADKQKNGIKV
NFKIRHNIEDGSVQLADHYQQNTPIGDGPVLLPDNHYLSTQSALSKDPNEKRDHMVLLEFV
TAAGITLGMDELYKSGLRSRQGSGAGSGAGSGAGSGAGSGAPRAQASNSAVDGTAGPG*-
[EMCV IRES]-MTEYKPTVRLATRDDVPRAVRTLAAAFADYPATRHTVDPDRHIERVTEL
QELFLTRVGLDIGKVWVADDGAAVAVWTTPESVEAGAVFAEIGPRMAELSGSRLAAQQQME
GLLAPHRPKEPAWFLATVGVSPDHQGKGLGSAVVLPGVEAAERAGVPAFLETSAPRNLPFY
ERLGFTVTADVECPKDRATWCMTRKPGA*
```

```
>DNA sequence
ATGAAAAAAAAGAAAAAGAAAGGCAGCGGTGCTGACAAAGACTGCGAAATGAAGCGCACCA
CCCTGGATAGCCCTCTGGGCAAGCTGGAACTGTCTGGGTGCGAACAGGGCCTGCACCGTAT
CATCTTCCTGGGCAAAGGAACATCTGCCGCCGACGCCGTGGAAGTGCCTGCCCCAGCCGCC
GTGCTGGGCGGACCAGAGCCACTGATGCAGGCCACCGCCTGGCTCAACGCCTACTTTCACC
AGCCTGAGGCCATCGAGGAGTTCCCTGTGCCAGCCCTGCACCACCCAGTGTTCCAGCAGGA
GAGCTTTACCCGCCAGGTGCTGTGGAAACTGCTGAAAGTGGTGAAGTTCGGAGAGGTCATC
AGCTACAGCCACCTGGCCGCCCTGGCCGGCAATCCCGCCGCCACCGCCGCCGTGAAAACCG
CCCTGAGCGGAAATCCCGTGCCCATTCTGATCCCCTGCCACCGGGTGGTGCAGGGCGACCT
GGACGTGGGGGGCTACGAGGGCGGGCTCGCCGTGAAAGAGTGGCTGCTGGCCCACGAGGGC
CACAGACTGGGCAAGCCTGGGCTGGGTGCGGCCGCTTCGGACCCACCGGTCGCCACCATGG
TGAGCAAGGGCGAGGAGCTGTTCACCGGGGTGGTGCCCATCCTGGTCGAGCTGGACGGCGA
CGTAAACGGCCACAAGTTCAGCGTGTCCGGCGAGGGCGAGGGCGATGCCACCTACGGCAAG
CTGACCCTGAAGTTCATCTGCACCACCGGCAAGCTGCCCGTGCCCTGGCCCACCCTCGTGA
CCACCCTGACCTACGGCGTGCAGTGCTTCAGCCGCTACCCCGACCACATGAAGCAGCACGA
CTTCTTCAAGTCCGCCATGCCCGAAGGCTACGTCCAGGAGCGCACCATCTTCTTCAAGGAC
GACGGCAACTACAAGACCCGCGCCGAGGTGAAGTTCGAGGGCGACACCCTGGTGAACCGCA
TCGAGCTGAAGGGCATCGACTTCAAGGAGGACGGCAACATCCTGGGGCACAAGCTGGAGTA
CAACTACAACAGCCACAACGTCTATATCATGGCCGACAAGCAGAAGAACGGCATCAAGGTG
AACTTCAAGATCCGCCACAACATCGAGGACGGCAGCGTGCAGCTCGCCGACCACTACCAGC
AGAACACCCCCATCGGCGACGGCCCCGTGCTGCTGCCCGACAACCACTACCTGAGCACCCA
GTCCGCCCTGAGCAAAGACCCCAACGAGAAGCGCGATCACATGGTCCTGCTGGAGTTCGTG
ACCGCCGCCGGGATCACTCTCGGCATGGACGAGCTGTACAAGTCCGGACTCAGATCTCGAC
AAGGTAGTGGTGCTGGCTCTGGTGCTGGTAGTGGCGCTGGTTCCGGTGCTGGCTCTGGCGC
GCCTCGAGCTCAAGCTTCGAATTCTGCAGTCGACGGTACCGCGGGCCCGGGATAAGTCAAC
TAACTTAAGCTAGCAACGGTTTCCCTCTAGCGGGATCAATTcccccccccctaacgttac
tggccgaagccgcttggaataaggccggtgtgcgtttgtctatatgttattttccaccata
ttgccgtcttttggcaatgtgagggcccggaaacctggccctgtcttcttgacgagcattc
ctaggggtctttcccctctcgccaaaggaatgcaaggtctgttgaatgtcgtgaaggaagc
agttcctctggaagcttcttgaagacaaacaacgtctgtagcgaccctttgcaggcagcgg
aaccccccacctggcgacaggtgcctctgcggccaaaagccacgtgtataagatacacctg
caaaggcggcacaaccccagtgccacgttgtgagttggatagttgtggaaagagtcaaatg
gctctcctcaagcgtattcaacaaggggctgaaggatgcccagaaggtaccccattgtatg
ggatctgatctggggcctcggtgcacatgctttacatgtgtttagtcgaggttaaaaaacg
tctaggccccccgaaccacggggacgtggttttcctttgaaaaacacgatAATACCATGAC
CGAGTACAAGCCCACGGTGCGCCTCGCCACCCGCGACGACGTCCCCAGGGCCGTACGCACC
```

CTCGCCGCCGCGTTCGCCGACTACCCCGCCACGCGCCACACCGTCGATCCGGACCGCCACA
TCGAGCGGGTCACCGAGCTGCAAGAACTCTTCCTCACGCGCGTCGGGCTCGACATCGGCAA
GGTGTGGGTCGCGGACGACGGCGCCGCGGTGGCGGTCTGGACCACGCCGGAGAGCGTCGAA
GCGGGGGCGGTGTTCGCCGAGATCGGCCCGCGCATGGCCGAGTTGAGCGGTTCCCGGCTGG
CCGCGCAGCAACAGATGGAAGGCCTCCTGGCGCCGCACCGGCCCAAGGAGCCCGCGTGGTT
CCTGGCCACCGTCGGCGTCTCGCCCGACCACCAGGGCAAGGGTCTGGGCAGCGCCGTCGTG
CTCCCCGGAGTGGAGGCGGCCGAGCGCGCCGGGGTGCCCGCCTTCCTGGAGACCTCCGCGC
CCCGCAACCTCCCCTTCTACGAGCGGCTCGGCTTCACCGTCACCGCCGACGTCGAGTGCCC
GAAGGACCGCGCGACCTGGTGCATGACCCGCAAGCCCGGTGCCTGA

**e**

pPBpuro-K6-SNAP$_i$-EGFP

>Amino acid sequence
MKKKKKKGSGADKDCEMKRTTLDSPLGKLELSGCEQGLHRIIFLGKGTSAADAVEVPAPAA
VLGGPEPLMQATAWLNAYFHQPEAIEEFPVPALHHPVFQQESFTRQVLWKLLKVVKFGEVI
SYSHLAALAGNPAATAAVKTALSGNPVPILIPCHRVVQGDLDVGGYEGGLAVKEWLLAHEA
AASDPPVATMVSKGEELFTGVVPILVELDGDVNGHKFSVSGEGEGDATYGKLTLKFICTTG
KLPVPWPTLVTTLTYGVQCFSRYPDHMKQHDFFKSAMPEGYVQERTIFFKDDGNYKTRAEV
KFEGDTLVNRIELKGIDFKEDGNILGHKLEYNYNSHNVYIMADKQKNGIKVNFKIRHNIED
GSVQLADHYQQNTPIGDGPVLLPDNHYLSTQSALSKDPNEKRDHMVLLEFVTAAGITLGMD
ELYKSGLRSRQGSGAGSGAGSGAGSGAGSGAPRAQASNSAVDGTAGPG*-[EMCV  IRES]
-MTEYKPTVRLATRDDVPRAVRTLAAAFADYPATRHTVDPDRHIERVTELQELFLTRVGLD
IGKVWVADDGAAVAVWTTPESVEAGAVFAEIGPRMAELSGSRLAAQQQMEGLLAPHRPKEP
AWFLATVGVSPDHQGKGLGSAVVLPGVEAAERAGVPAFLETSAPRNLPFYERLGFTVTADV
ECPKDRATWCMTRKPGA*

>DNA sequence
ATGAAAAAAAAGAAAAAGAAAGGCAGCGGTGCTGACAAAGACTGCGAAATGAAGCGCACCA
CCCTGGATAGCCCTCTGGGCAAGCTGGAACTGTCTGGGTGCGAACAGGGCCTGCACCGTAT
CATCTTCCTGGGCAAAGGAACATCTGCCGCCGACGCCGTGGAAGTGCCTGCCCCAGCCGCC
GTGCTGGGCGGACCAGAGCCACTGATGCAGGCCACCGCCTGGCTCAACGCCTACTTTCACC
AGCCTGAGGCCATCGAGGAGTTCCCTGTGCCAGCCCTGCACCACCCAGTGTTCCAGCAGGA
GAGCTTTACCCGCCAGGTGCTGTGGAAACTGCTGAAAGTGGTGAAGTTCGGAGAGGTCATC
AGCTACAGCCACCTGGCCGCCCTGGCCGGCAATCCCGCCGCCACCGCCGCCGTGAAAACCG
CCCTGAGCGGAAATCCCGTGCCCATTCTGATCCCCTGCCACCGGGTGGTGCAGGGCGACCT
GGACGTGGGGGGCTACGAGGGCGGGCTCGCCGTGAAAGAGTGGCTGCTGGCCCACGAGGCG
GCCGCTTCGGACCCACCGGTCGCCACCATGGTGAGCAAGGGCGAGGAGCTGTTCACCGGGG
TGGTGCCCATCCTGGTCGAGCTGGACGGCGACGTAAACGGCCACAAGTTCAGCGTGTCCGG
CGAGGGCGAGGGCGATGCCACCTACGGCAAGCTGACCCTGAAGTTCATCTGCACCACCGGC
AAGCTGCCCGTGCCCTGGCCCACCCTCGTGACCACCCTGACCTACGGCGTGCAGTGCTTCA
GCCGCTACCCCGACCACATGAAGCAGCACGACTTCTTCAAGTCCGCCATGCCCGAAGGCTA
CGTCCAGGAGCGCACCATCTTCTTCAAGGACGACGGCAACTACAAGACCCGCGCCGAGGTG
AAGTTCGAGGGCGACACCCTGGTGAACCGCATCGAGCTGAAGGGCATCGACTTCAAGGAGG
ACGGCAACATCCTGGGGCACAAGCTGGAGTACAACTACAACAGCCACAACGTCTATATCAT
GGCCGACAAGCAGAAGAACGGCATCAAGGTGAACTTCAAGATCCGCCACAACATCGAGGAC
GGCAGCGTGCAGCTCGCCGACCACTACCAGCAGAACACCCCCATCGGCGACGGCCCCGTGC
TGCTGCCCGACAACCACTACCTGAGCACCCAGTCCGCCCTGAGCAAAGACCCCAACGAGAA
GCGCGATCACATGGTCCTGCTGGAGTTCGTGACCGCCGCCGGGATCACTCTCGGCATGGAC
GAGCTGTACAAGTCCGGACTCAGATCTCGACAAGGTAGTGGTGCTGGCTCTGGTGCTGGTA
GTGGCGCTGGTTCCGGTGCTGGCTCTGGCGCGCCTCGAGCTCAAGCTTCGAATTCTGCAGT
CGACGGTACCGCGGGCCCGGGATAAGTCAACTAACTTAAGCTAGCAACGGTTTCCCTCTAG
CGGGATCAATTcccccccccctaacgttactggccgaagccgcttggaataaggccggtg
tgcgtttgtctatatgttattttccaccatattgccgtcttttggcaatgtgagggcccgg
aaacctggccctgtcttcttgacgagcattcctaggggtctttcccctctcgccaaaggaa
tgcaaggtctgttgaatgtcgtgaaggaagcagttcctctggaagcttcttgaagacaaac
aacgtctgtagcgaccctttgcaggcagcggaaccccccacctggcgacaggtgcctctgc
ggccaaaagccacgtgtataagatacacctgcaaaggcggcacaaccccagtgccacgttg
tgagttggatagttgtggaaagagtcaaatggctctcctcaagcgtattcaacaaggggct
gaaggatgcccagaaggtaccccattgtatgggatctgatctggggcctcggtgcacatgc
tttacatgtgtttagtcgaggttaaaaaacgtctaggccccccgaaccacggggacgtggt
tttcctttgaaaaacacgatAATACCATGACCGAGTACAAGCCCACGGTGCGCCTCGCCAC
CCGCGACGACGTCCCCAGGGCCGTACGCACCCTCGCCGCCGCGTTCGCCGACTACCCCGCC

ACGCGCCACACCGTCGATCCGGACCGCCACATCGAGCGGGTCACCGAGCTGCAAGAACTCT
TCCTCACGCGCGTCGGGCTCGACATCGGCAAGGTGTGGGTCGCGGACGACGGCGCCGCGGT
GGCGGTCTGGACCACGCCGGAGAGCGTCGAAGCGGGGGCGGTGTTCGCCGAGATCGGCCCG
CGCATGGCCGAGTTGAGCGGTTCCCGGCTGGCCGCGCAGCAACAGATGGAAGGCCTCCTGG
CGCCGCACCGGCCCAAGGAGCCCGCGTGGTTCCTGGCCACCGTCGGCGTCTCGCCCGACCA
CCAGGGCAAGGGTCTGGGCAGCGCCGTCGTGCTCCCCGGAGTGGAGGCGGCCGAGCGCGCC
GGGGTGCCCGCCTTCCTGGAGACCTCCGCGCCCCGCAACCTCCCCTTCTACGAGCGGCTCG
GCTTCACCGTCACCGCCGACGTCGAGTGCCCGAAGGACCGCGCGACCTGGTGCATGACCCG
CAAGCCCGGTGCCTGA

**f**

pPBpuro-K6-SNAP$_f$-EGFP-Tiam1

>Amino acid sequence
MKKKKKKGSGADKDCEMKRTTLDSPLGKLELSGCEQGLHRIIFLGKGTSAADAVEVPAPAA
VLGGPEPLMQATAWLNAYFHQPEAIEEFPVPALHHPVFQQESFTRQVLWKLLKVVKFGEVI
SYSHLAALAGNPAATAAVKTALSGNPVPILIPCHRVVQGDLDVGGYEGGLAVKEWLLAHEG
HRLGKPGLGAAASDPPVATMVSKGEELFTGVVPILVELDGDVNGHKFSVSGEGEGDATYGK
LTLKFICTTGKLPVPWPTLVTTLTYGVQCFSRYPDHMKQHDFFKSAMPEGYVQERTIFFKD
DGNYKTRAEVKFEGDTLVNRIELKGIDFKEDGNILGHKLEYNYNSHNVYIMADKQKNGIKV
NFKIRHNIEDGSVQLADHYQQNTPIGDGPVLLPDNHYLSTQSALSKDPNEKRDHMVLLEFV
TAAGITLGMDELYKSGLRSRQGSGAGSGAGSGAGSGAGSGAPRAMNPSDQNPSPQDSTGPQ
LATMRQLSDADNVRKVICELLETERTYVKDLNCLMERYLKPLQKETFLTQDELDVLFGNLT
EMVEFQVEFLKTLEDGVRLVPDLEKLEKVDQFKKVLFSLGGSFLYYADRFKLYSAFCAIHT
KVPKVLVKAKTDTAFKAFLDAQNPKQQHSSTLESYLIKPIQRILKYPLLLRELFALTDAES
EEHYHLDVAIKTMNKVASHINEMQKIHEEFGAVFDQLIAEQTGEKKEVADLSMGDLLLHTT
VIWLNPPASLGKWKKEPELAAFVFKTAVVLVYKDGSKQKKKLVGSHRLSIYEDWDPFRFRH
MIPTEALQVRALASADAEANAVCEIVHVKSESEGRPERVFHLCCSSPESRKDFLKAVHSIL
RDKHRRQLLKTESLPSSQQYVPFGGKRLCALKGARPAMSRAVSAPSKSLGRRRRRLARNRF
TIDSDAVSASSPEKESQQPPGGGDTDRWVEEQFDLAQYEEQDDIKETDILSDDDEFCESVK
GASVDRDLQERLQATSISQRERGRKTLDSHASRMAQLKKQAALSGINGGLESASEEVIWVR
REDFAPSRKLNTEI*

>DNA sequence
ATGAAAAAAAAGAAAAAGAAAGGCAGCGGTGCTGACAAAGACTGCGAAATGAAGCGCACCA
CCCTGGATAGCCCTCTGGGCAAGCTGGAACTGTCTGGGTGCGAACAGGGCCTGCACCGTAT
CATCTTCCTGGGCAAAGGAACATCTGCCGCCGACGCCGTGGAAGTGCCTGCCCCAGCCGCC
GTGCTGGGCGGACCAGAGCCACTGATGCAGGCCACCGCCTGGCTCAACGCCTACTTTCACC
AGCCTGAGGCCATCGAGGAGTTCCCTGTGCCAGCCCTGCACCACCCAGTGTTCCAGCAGGA
GAGCTTTACCCGCCAGGTGCTGTGGAAACTGCTGAAAGTGGTGAAGTTCGGAGAGGTCATC
AGCTACAGCCACCTGGCCGCCCTGGCCGGCAATCCCGCCGCCACCGCCGCCGTGAAAACCG
CCCTGAGCGGAAATCCCGTGCCCATTCTGATCCCCTGCCACCGGGTGGTGCAGGGCGACCT
GGACGTGGGGGGCTACGAGGGCGGGCTCGCCGTGAAAGAGTGGCTGCTGGCCCACGAGGGC
CACAGACTGGGCAAGCCTGGGCTGGGTGCGGCCGCTTCGGACCCACCGGTCGCCACCATGG
TGAGCAAGGGCGAGGAGCTGTTCACCGGGGTGGTGCCCATCCTGGTCGAGCTGGACGGCGA
CGTAAACGGCCACAAGTTCAGCGTGTCCGGCGAGGGCGAGGGCGATGCCACCTACGGCAAG
CTGACCCTGAAGTTCATCTGCACCACCGGCAAGCTGCCCGTGCCCTGGCCCACCCTCGTGA
CCACCCTGACCTACGGCGTGCAGTGCTTCAGCCGCTACCCCGACCACATGAAGCAGCACGA
CTTCTTCAAGTCCGCCATGCCCGAAGGCTACGTCCAGGAGCGCACCATCTTCTTCAAGGAC
GACGGCAACTACAAGACCCGCGCCGAGGTGAAGTTCGAGGGCGACACCCTGGTGAACCGCA
TCGAGCTGAAGGGCATCGACTTCAAGGAGGACGGCAACATCCTGGGGCACAAGCTGGAGTA
CAACTACAACAGCCACAACGTCTATATCATGGCCGACAAGCAGAAGAACGGCATCAAGGTG
AACTTCAAGATCCGCCACAACATCGAGGACGGCAGCGTGCAGCTCGCCGACCACTACCAGC
AGAACACCCCCATCGGCGACGGCCCCGTGCTGCTGCCCGACAACCACTACCTGAGCACCCA
GTCCGCCCTGAGCAAAGACCCCAACGAGAAGCGCGATCACATGGTCCTGCTGGAGTTCGTG
ACCGCCGCCGGGATCACTCTCGGCATGGACGAGCTGTACAAGTCCGGACTCAGATCTCGAC
AAGGTAGTGGTGCTGGCTCTGGTGCTGGTAGTGGCGCTGGTTCCGGTGCTGGCTCTGGCGC
GCCTCGAGCAATGAACCCCTCTGACCAGAACCCATCTCCTCAGGACTCCACGGGGCCTCAG
CTGGCGACCATGAGACAACTCTCGGATGCAGATAACGTGCGCAAGGTGATCTGCGAGCTCC
TGGAGACGGAGCGCACCTACGTGAAGGATTTAAACTGTCTTATGGAGAGATACCTAAAGCC
TCTTCAAAAAGAAACTTTTCTCACCCAGGATGAGCTTGACGTGCTTTTTGGAAATTTAACG
GAAATGGTAGAGTTTCAAGTAGAATTCCTTAAAACTCTAGAAGATGGAGTGAGACTGGTAC
CTGATTTGGAAAAGCTTGAGAAGGTTGATCAATTTAAGAAAGTGCTGTTCTCTCTGGGGGG

ATCATTCCTGTATTATGCTGACCGCTTCAAGCTCTACAGTGCCTTCTGCGCCATCCACACA
AAAGTTCCCAAGGTCCTGGTGAAAGCCAAGACAGACACGGCTTTCAAGGCATTCTTGGATG
CCCAGAACCCGAAGCAGCAGCACTCATCCACGCTGGAGTCGTACCTCATCAAGCCCATCCA
GAGGATCCTCAAGTACCCACTTCTGCTCAGGGAGCTGTTCGCCCTGACCGATGCGGAGAGC
GAGGAGCACTACCACCTGGACGTGGCCATCAAGACCATGAACAAGGTTGCCAGTCACATCA
ATGAGATGCAGAAAATCCATGAAGAGTTTGGGGCTGTGTTTGACCAGCTGATTGCTGAACA
GACTGGTGAGAAAAAAGAGGTTGCAGATCTGAGCATGGGAGACCTGCTTTTGCACACTACC
GTGATCTGGCTGAACCCGCCGGCCTCGCTGGGCAAGTGGAAAAAGGAACCAGAGTTGGCAG
CATTCGTCTTCAAAACTGCTGTGGTCCTTGTGTATAAAGATGGTTCCAAACAGAAGAAGAA
ACTTGTAGGATCTCACAGGCTTTCCATTTATGAGGACTGGGACCCCTTCAGATTTCGACAC
ATGATCCCCACGGAAGCGCTGCAGGTTCGAGCTTTGGCGAGTGCAGATGCAGAGGCAAATG
CCGTGTGTGAAATTGTCCATGTAAAATCCGAGTCTGAAGGGAGGCCGGAGAGGGTCTTTCA
CTTGTGCTGCAGCTCCCCAGAGAGCCGAAAGGATTTCCTAAAGGCTGTGCATTCAATCCTG
CGTGATAAGCACAGAAGACAGCTCCTCAAAACCGAGAGCCTTCCCTCATCCCAGCAATATG
TCCCTTTTGGAGGCAAAAGATTGTGTGCACTGAAGGGGGCCAGGCCGGCCATGAGCAGGGC
AGTGTCTGCCCCAAGCAAGTCTCTTGGGAGGAGGAGGCGGCGGCTGGCTCGAAACAGGTTT
ACCATTGATTCTGATGCCGTCTCCGCAAGCAGCCCGGAGAAAGAGTCCCAGCAGCCCCCCG
GTGGTGGGGACACTGACCGATGGGTAGAGGAGCAGTTTGATCTTGCTCAGTATGAGGAGCA
AGATGACATCAAGGAGACAGACATCCTCAGTGACGATGATGAGTTCTGTGAGTCCGTGAAG
GGTGCCTCAGTGGACAGAGACCTGCAGGAGCGGCTTCAGGCCACCTCCATCAGTCAGCGGG
AAAGAGGCCGGAAAACCCTGGATAGTCACGCGTCCCGCATGGCACAGCTCAAGAAGCAAGC
TGCCCTGTCGGGGATCAATGGAGGCCTGGAGAGCGCAAGCGAGGAAGTCATTTGGGTTAGG
CGTGAAGACTTTGCCCCCTCCAGGAAACTGAACACTGAGATCTGA

**g**

pPBpuro-K6-SNAP$_i$-EGFP-Tiam1

>Amino acid sequence
MKKKKKKGSGADKDCEMKRTTLDSPLGKLELSGCEQGLHRIIFLGKGTSAADAVEVPAPAA
VLGGPEPLMQATAWLNAYFHQPEAIEEFPVPALHHPVFQQESFTRQVLWKLLKVVKFGEVI
SYSHLAALAGNPAATAAVKTALSGNPVPILIPCHRVVQGDLDVGGYEGGLAVKEWLLAHEA
AASDPPVATMVSKGEELFTGVVPILVELDGDVNGHKFSVSGEGEGDATYGKLTLKFICTTG
KLPVPWPTLVTTLTYGVQCFSRYPDHMKQHDFFKSAMPEGYVQERTIFFKDDGNYKTRAEV
KFEGDTLVNRIELKGIDFKEDGNILGHKLEYNYNSHNVYIMADKQKNGIKVNFKIRHNIED
GSVQLADHYQQNTPIGDGPVLLPDNHYLSTQSALSKDPNEKRDHMVLLEFVTAAGITLGMD
ELYKSGLRSRQGSGAGSGAGSGAGSGAGSGAPRAMNPSDQNPSPQDSTGPQLATMRQLSDA
DNVRKVICELLETERTYVKDLNCLMERYLKPLQKETFLTQDELDVLFGNLTEMVEFQVEFL
KTLEDGVRLVPDLEKLEKVDQFKKVLFSLGGSFLYYADRFKLYSAFCAIHTKVPKVLVKAK
TDTAFKAFLDAQNPKQQHSSTLESYLIKPIQRILKYPLLLRELFALTDAESEEHYHLDVAI
KTMNKVASHINEMQKIHEEFGAVFDQLIAEQTGEKKEVADLSMGDLLLHTTVIWLNPPASL
GKWKKEPELAAFVFKTAVVLVYKDGSKQKKKLVGSHRLSIYEDWDPFRFRHMIPTEALQVR
ALASADAEANAVCEIVHVKSESEGRPERVFHLCCSSPESRKDFLKAVHSILRDKHRRQLLK
TESLPSSQQYVPFGGKRLCALKGARPAMSRAVSAPSKSLGRRRRRLARNRFTIDSDAVSAS
SPEKESQQPPGGGDTDRWVEEQFDLAQYEEQDDIKETDILSDDDEFCESVKGASVDRDLQE
RLQATSISQRERGRKTLDSHASRMAQLKKQAALSGINGGLESASEEVIWVRREDFAPSRKL
NTEI*

>DNA sequence
ATGAAAAAAAAGAAAAAGAAAGGCAGCGGTGCTGACAAAGACTGCGAAATGAAGCGCACCA
CCCTGGATAGCCCTCTGGGCAAGCTGGAACTGTCTGGGTGCGAACAGGGCCTGCACCGTAT
CATCTTCCTGGGCAAAGGAACATCTGCCGCCGACGCCGTGGAAGTGCCTGCCCCAGCCGCC
GTGCTGGGCGGACCAGAGCCACTGATGCAGGCCACCGCCTGGCTCAACGCCTACTTTCACC
AGCCTGAGGCCATCGAGGAGTTCCCTGTGCCAGCCCTGCACCACCCAGTGTTCCAGCAGGA
GAGCTTTACCCGCCAGGTGCTGTGGAAACTGCTGAAAGTGGTGAAGTTCGGAGAGGTCATC
AGCTACAGCCACCTGGCCGCCCTGGCCGGCAATCCCGCCGCCACCGCCGCCGTGAAAACCG
CCCTGAGCGGAAATCCCGTGCCCATTCTGATCCCCTGCCACCGGGTGGTGCAGGGCGACCT
GGACGTGGGGGGCTACGAGGGCGGGCTCGCCGTGAAAGAGTGGCTGCTGGCCCACGAGGCG
GCCGCTTCGGACCCACCGGTCGCCACCATGGTGAGCAAGGGCGAGGAGCTGTTCACCGGGG
TGGTGCCCATCCTGGTCGAGCTGGACGGCGACGTAAACGGCCACAAGTTCAGCGTGTCCGG
CGAGGGCGAGGGCGATGCCACCTACGGCAAGCTGACCCTGAAGTTCATCTGCACCACCGGC
AAGCTGCCCGTGCCCTGGCCCACCCTCGTGACCACCCTGACCTACGGCGTGCAGTGCTTCA
GCCGCTACCCCGACCACATGAAGCAGCACGACTTCTTCAAGTCCGCCATGCCCGAAGGCTA
CGTCCAGGAGCGCACCATCTTCTTCAAGGACGACGGCAACTACAAGACCCGCGCCGAGGTG
AAGTTCGAGGGCGACACCCTGGTGAACCGCATCGAGCTGAAGGGCATCGACTTCAAGGAGG
ACGGCAACATCCTGGGGCACAAGCTGGAGTACAACTACAACAGCCACAACGTCTATATCAT
GGCCGACAAGCAGAAGAACGGCATCAAGGTGAACTTCAAGATCCGCCACAACATCGAGGAC
GGCAGCGTGCAGCTCGCCGACCACTACCAGCAGAACACCCCCATCGGCGACGGCCCCGTGC
TGCTGCCCGACAACCACTACCTGAGCACCCAGTCCGCCCTGAGCAAAGACCCCAACGAGAA
GCGCGATCACATGGTCCTGCTGGAGTTCGTGACCGCCGCCGGGATCACTCTCGGCATGGAC
GAGCTGTACAAGTCCGGACTCAGATCTCGACAAGGTAGTGGTGCTGGCTCTGGTGCTGGTA
GTGGCGCTGGTTCCGGTGCTGGCTCTGGCGCGCCTCGAGCAATGAACCCCTCTGACCAGAA
CCCATCTCCTCAGGACTCCACGGGGCCTCAGCTGGCGACCATGAGACAACTCTCGGATGCA
GATAACGTGCGCAAGGTGATCTGCGAGCTCCTGGAGACGGAGCGCACCTACGTGAAGGATT
TAAACTGTCTTATGGAGAGATACCTAAAGCCTCTTCAAAAAGAAACTTTTCTCACCCAGGA
TGAGCTTGACGTGCTTTTTGGAAATTTAACGGAAATGGTAGAGTTTCAAGTAGAATTCCTT
AAAACTCTAGAAGATGGAGTGAGACTGGTACCTGATTTGGAAAAGCTTGAGAAGGTTGATC
AATTTAAGAAAGTGCTGTTCTCTCTGGGGGGATCATTCCTGTATTATGCTGACCGCTTCAA

GCTCTACAGTGCCTTCTGCGCCATCCACACAAAAGTTCCCAAGGTCCTGGTGAAAGCCAAG
ACAGACACGGCTTTCAAGGCATTCTTGGATGCCCAGAACCCGAAGCAGCAGCACTCATCCA
CGCTGGAGTCGTACCTCATCAAGCCCATCCAGAGGATCCTCAAGTACCCACTTCTGCTCAG
GGAGCTGTTCGCCCTGACCGATGCGGAGAGCGAGGAGCACTACCACCTGGACGTGGCCATC
AAGACCATGAACAAGGTTGCCAGTCACATCAATGAGATGCAGAAAATCCATGAAGAGTTTG
GGGCTGTGTTTGACCAGCTGATTGCTGAACAGACTGGTGAGAAAAAAGAGGTTGCAGATCT
GAGCATGGGAGACCTGCTTTTGCACACTACCGTGATCTGGCTGAACCCGCCGGCCTCGCTG
GGCAAGTGGAAAAAGGAACCAGAGTTGGCAGCATTCGTCTTCAAAACTGCTGTGGTCCTTG
TGTATAAAGATGGTTCCAAACAGAAGAAGAAACTTGTAGGATCTCACAGGCTTTCCATTTA
TGAGGACTGGGACCCCTTCAGATTTCGACACATGATCCCCACGGAAGCGCTGCAGGTTCGA
GCTTTGGCGAGTGCAGATGCAGAGGCAAATGCCGTGTGTGAAATTGTCCATGTAAAATCCG
AGTCTGAAGGGAGGCCGGAGAGGGTCTTTCACTTGTGCTGCAGCTCCCCAGAGAGCCGAAA
GGATTTCCTAAAGGCTGTGCATTCAATCCTGCGTGATAAGCACAGAAGACAGCTCCTCAAA
ACCGAGAGCCTTCCCTCATCCCAGCAATATGTCCCTTTTGGAGGCAAAAGATTGTGTGCAC
TGAAGGGGGCCAGGCCGGCCATGAGCAGGGCAGTGTCTGCCCCAAGCAAGTCTCTTGGGAG
GAGGAGGCGGCGGCTGGCTCGAAACAGGTTTACCATTGATTCTGATGCCGTCTCCGCAAGC
AGCCCGGAGAAAGAGTCCCAGCAGCCCCCCGGTGGTGGGGACACTGACCGATGGGTAGAGG
AGCAGTTTGATCTTGCTCAGTATGAGGAGCAAGATGACATCAAGGAGACAGACATCCTCAG
TGACGATGATGAGTTCTGTGAGTCCGTGAAGGGTGCCTCAGTGGACAGAGACCTGCAGGAG
CGGCTTCAGGCCACCTCCATCAGTCAGCGGGAAAGAGGCCGGAAAACCCTGGATAGTCACG
CGTCCCGCATGGCACAGCTCAAGAAGCAAGCTGCCCTGTCGGGGATCAATGGAGGCCTGGA
GAGCGCAAGCGAGGAAGTCATTTGGGTTAGGCGTGAAGACTTTGCCCCCTCCAGGAAACTG
AACACTGAGATCTGA

**h**

pCMV-Lifeact-mCherry

>Amino acid sequence
MGVADLIKKFESISKEEGDPPVATMVSKGEEDNMAIIKEFMRFKVHMEGSVNGHEFEIEGE
GEGRPYEGTQTAKLKVTKGGPLPFAWDILSPQFMYGSKAYVKHPADIPDYLKLSFPEGFKW
ERVMNFEDGGVVTVTQDSSLQDGEFIYKVKLRGTNFPSDGPVMQKKTMGWEASSERMYPED
GALKGEIKQRLKLKDGGHYDAEVKTTYKAKKPVQLPGAYNVNIKLDITSHNEDYTIVEQYE
RAEGRHSTGGMDELYK*

>DNA sequence
ATGGGCGTGGCCGACTTGATCAAGAAGTTCGAGTCCATCTCCAAGGAGGAGGGGGATCCAC
CGGTCGCCACCATGGTGAGCAAGGGCGAGGAGGATAACATGGCCATCATCAAGGAGTTCAT
GCGCTTCAAGGTGCACATGGAGGGCTCCGTGAACGGCCACGAGTTCGAGATCGAGGGCGAG
GGCGAGGGCCGCCCCTACGAGGGCACCCAGACCGCCAAGCTGAAGGTGACCAAGGGTGGCC
CCCTGCCCTTCGCCTGGGACATCCTGTCCCCTCAGTTCATGTACGGCTCCAAGGCCTACGT
GAAGCACCCCGCCGACATCCCCGACTACTTGAAGCTGTCCTTCCCCGAGGGCTTCAAGTGG
GAGCGCGTGATGAACTTCGAGGACGGCGGCGTGGTGACCGTGACCCAGGACTCCTCCCTGC
AGGACGGCGAGTTCATCTACAAGGTGAAGCTGCGCGGCACCAACTTCCCCTCCGACGGCCC
CGTAATGCAGAAGAAGACCATGGGCTGGGAGGCCTCCTCCGAGCGGATGTACCCCGAGGAC
GGCGCCCTGAAGGGCGAGATCAAGCAGAGGCTGAAGCTGAAGGACGGCGGCCACTACGACG
CTGAGGTCAAGACCACCTACAAGGCCAAGAAGCCCGTGCAGCTGCCCGGCGCCTACAACGT
CAACATCAAGTTGGACATCACCTCCCACAACGAGGACTACACCATCGTGGAACAGTACGAA
CGCGCCGAGGGCCGCCACTCCACCGGCGGCATGGACGAGCTGTACAAGTAG

**i**

pPBpuro-K6-SNAP$_i$-EGFP-cRaf

>Amino acid sequence

MKKKKKKGSGADKDCEMKRTTLDSPLGKLELSGCEQGLHRIIFLGKGTSAADAVEVPAPAA
VLGGPEPLMQATAWLNAYFHQPEAIEEFPVPALHHPVFQQESFTRQVLWKLLKVVKFGEVI
SYSHLAALAGNPAATAAVKTALSGNPVPILIPCHRVVQGDLDVGGYEGGLAVKEWLLAHEA
AASDPPVATMVSKGEELFTGVVPILVELDGDVNGHKFSVSGEGEGDATYGKLTLKFICTTG
KLPVPWPTLVTTLTYGVQCFSRYPDHMKQHDFFKSAMPEGYVQERTIFFKDDGNYKTRAEV
KFEGDTLVNRIELKGIDFKEDGNILGHKLEYNYNSHNVYIMADKQKNGIKVNFKIRHNIED
GSVQLADHYQQNTPIGDGPVLLPDNHYLSTQSALSKDPNEKRDHMVLLEFVTAAGITLGMD
ELYKSGLRSRQGSGAGSGAGSGAGSGAGSGAPRSLEMEHIQGAWKTISNGFGFKDAVFDGS
SCISPTIVQQFGYQRRASDDGKLTDPSKTSNTIRVFLPNKQRTVVNVRNGMSLHDCLMKAL
KVRGLQPECCAVFRLLHEHKGKKARLDWNTDAASLIGEELQVDFLDHVPLTTHNFARKTFL
KLAFCDICQKFLLNGFRCQTCGYKFHEHCSTKVPTMCVDWSNIRQLLLFPNSTIGDSGVPA
LPSLTMRRMRESVSRMPVSSQHRYSTPHAFTFNTSSPSSEGSLSQRQRSTSTPNVHMVSTT
LPVDSRMIEDAIRSHSESASPSALSSSPNNLSPTGWSQPKTPVPAQRERAPVSGTQEKNKI
RPRGQRDSSYYWEIEASEVMLSTRIGSGSFGTVYKGKWHGDVAVKILKVVDPTPEQFQAFR
NEVAVLRKTRHVNILLFMGYMTKDNLAIVTQWCEGSSLYKHLHVQETKFQMFQLIDIARQT
AQGMDYLHAKNIIHRDMKSNNIFLHEGLTVKIGDFGLATVKSRWSGSQQVEQPTGSVLWMA
PEVIRMQDNNPFSFQSDVYSYGIVLYELMTGELPYSHINNRDQIIFMVGRGYASPDLSKLY
KNCPKAMKRLVADCVKKVKEERPLFPQILSSIELLQHSLPKINRSASEPSLHRAAHTEDIN
ACTLTTSPRLPVFCGR*-[EMCV IRES]-MTEYKPTVRLATRDDVPRAVRTLAAAFADYP
ATRHTVDPDRHIERVTELQELFLTRVGLDIGKVWVADDGAAVAVWTTPESVEAGAVFAEIG
PRMAELSGSRLAAQQQMEGLLAPHRPKEPAWFLATVGVSPDHQGKGLGSAVVLPGVEAAER
AGVPAFLETSAPRNLPFYERLGFTVTADVECPKDRATWCMTRKPGA*

>DNA sequence

ATGAAAAAAAAGAAAAAGAAAGGCAGCGGTGCTGACAAAGACTGCGAAATGAAGCGCACCA
CCCTGGATAGCCCTCTGGGCAAGCTGGAACTGTCTGGGTGCGAACAGGGCCTGCACCGTAT
CATCTTCCTGGGCAAAGGAACATCTGCCGCCGACGCCGTGGAAGTGCCTGCCCCAGCCGCC
GTGCTGGGCGGACCAGAGCCACTGATGCAGGCCACCGCCTGGCTCAACGCCTACTTTCACC
AGCCTGAGGCCATCGAGGAGTTCCCTGTGCCAGCCCTGCACCACCCAGTGTTCCAGCAGGA
GAGCTTTACCCGCCAGGTGCTGTGGAAACTGCTGAAAGTGGTGAAGTTCGGAGAGGTCATC
AGCTACAGCCACCTGGCCGCCCTGGCCGGCAATCCCGCCGCCACCGCCGCCGTGAAAACCG
CCCTGAGCGGAAATCCCGTGCCCATTCTGATCCCCTGCCACCGGGTGGTGCAGGGCGACCT
GGACGTGGGGGGCTACGAGGGCGGGCTCGCCGTGAAAGAGTGGCTGCTGGCCCACGAGGCG
GCCGCTTCGGACCCACCGGTCGCCACCATGGTGAGCAAGGGCGAGGAGCTGTTCACCGGGG
TGGTGCCCATCCTGGTCGAGCTGGACGGCGACGTAAACGGCCACAAGTTCAGCGTGTCCGG
CGAGGGCGAGGGCGATGCCACCTACGGCAAGCTGACCCTGAAGTTCATCTGCACCACCGGC
AAGCTGCCCGTGCCCTGGCCCACCCTCGTGACCACCCTGACCTACGGCGTGCAGTGCTTCA
GCCGCTACCCCGACCACATGAAGCAGCACGACTTCTTCAAGTCCGCCATGCCCGAAGGCTA
CGTCCAGGAGCGCACCATCTTCTTCAAGGACGACGGCAACTACAAGACCCGCGCCGAGGTG
AAGTTCGAGGGCGACACCCTGGTGAACCGCATCGAGCTGAAGGGCATCGACTTCAAGGAGG
ACGGCAACATCCTGGGGCACAAGCTGGAGTACAACTACAACAGCCACAACGTCTATATCAT
GGCCGACAAGCAGAAGAACGGCATCAAGGTGAACTTCAAGATCCGCCACAACATCGAGGAC
GGCAGCGTGCAGCTCGCCGACCACTACCAGCAGAACACCCCCATCGGCGACGGCCCCGTGC
TGCTGCCCGACAACCACTACCTGAGCACCCAGTCCGCCCTGAGCAAAGACCCCAACGAGAA
GCGCGATCACATGGTCCTGCTGGAGTTCGTGACCGCCGCCGGGATCACTCTCGGCATGGAC
GAGCTGTACAAGTCCGGACTCAGATCTCGACAAGGTAGTGGTGCTGGCTCTGGTGCTGGTA
GTGGCGCTGGTTCCGGTGCTGGCTCTGGCGCGCCTCGAAGCCTCGAGATGGAGCACATACA
GGGAGCTTGGAAGACGATCAGCAATGGTTTTGGATTCAAAGATGCCGTGTTTGATGGCTCC
AGCTGCATCTCTCCTACAATAGTTCAGCAGTTTGGCTATCAGCGCCGGGCATCAGATGATG

GCAAACTCACAGATCCTTCTAAGACAAGCAACACTATCCGTGTTTTCTTGCCGAACAAGCA
AAGAACAGTGGTCAATGTGCGAAATGGAATGAGCTTGCATGACTGCCTTATGAAAGCACTC
AAGGTGAGGGGCCTGCAACCAGAGTGCTGTGCAGTGTTCAGACTTCTCCACGAACACAAAG
GTAAAAAAGCACGCTTAGATTGGAATACTGATGCTGCGTCTTTGATTGGAGAAGAACTTCA
AGTAGATTTCCTGGATCATGTTCCCCTCACAACACACAACTTTGCTCGGAAGACGTTCCTG
AAGCTTGCCTTCTGTGACATCTGTCAGAAATTCCTGCTCAATGGATTTCGATGTCAGACTT
GTGGCTACAAATTTCATGAGCACTGTAGCACCAAAGTACCTACTATGTGTGTGGACTGGAG
TAACATCAGACAACTCTTATTGTTTCCAAATTCCACTATTGGTGATAGTGGAGTCCCAGCA
CTACCTTCTTTGACTATGCGTCGTATGCGAGAGTCTGTTTCCAGGATGCCTGTTAGTTCTC
AGCACAGATATTCTACACCTCACGCCTTCACCTTTAACACCTCCAGTCCCTCATCTGAAGG
TTCCCTCTCCCAGAGGCAGAGGTCGACATCCACACCTAATGTCCACATGGTCAGCACCACG
CTGCCTGTGGACAGCAGGATGATTGAGGATGCAATTCGAAGTCACAGCGAATCAGCCTCAC
CTTCAGCCCTGTCCAGTAGCCCCAACAATCTGAGCCCAACAGGCTGGTCACAGCCGAAAAC
CCCCGTGCCAGCACAAAGAGAGCGGGCACCAGTATCTGGGACCCAGGAGAAAAACAAAATT
AGGCCTCGTGGACAGAGAGATTCAAGCTATTATTGGGAAATAGAAGCCAGTGAAGTGATGC
TGTCCACTCGGATTGGGTCAGGCTCTTTTGGAACTGTTTATAAGGGTAAATGGCACGGAGA
TGTTGCAGTAAAGATCCTAAAGGTTGTCGACCCAACCCCAGAGCAATTCCAGGCCTTCAGG
AATGAGGTGGCTGTTCTGCGCAAAACACGGCATGTGAACATTCTGCTTTTCATGGGGTACA
TGACAAAGGACAACCTGGCAATTGTGACCCAGTGGTGCGAGGGCAGCAGCCTCTACAAACA
CCTGCATGTCCAGGAGACCAAGTTTCAGATGTTCCAGCTAATTGACATTGCCCGGCAGACG
GCTCAGGGAATGGACTATTTGCATGCAAAGAACATCATCCATAGAGACATGAAATCCAACA
ATATATTTCTCCATGAAGGCTTAACAGTGAAAATTGGAGATTTTGGTTTGGCAACAGTAAA
GTCACGCTGGAGTGGTTCTCAGCAGGTTGAACAACCTACTGGCTCTGTCCTCTGGATGGCC
CCAGAGGTGATCCGAATGCAGGATAACAACCCATTCAGTTTCCAGTCGGATGTCTACTCCT
ATGGCATCGTATTGTATGAACTGATGACGGGGGAGCTTCCTTATTCTCACATCAACAACCG
AGATCAGATCATCTTCATGGTGGGCCGAGGATATGCCTCCCCAGATCTTAGTAAGCTATAT
AAGAACTGCCCCAAAGCAATGAAGAGGCTGGTAGCTGACTGTGTGAAGAAAGTAAAGGAAG
AGAGGCCTCTTTTTCCCCAGATCCTGTCTTCCATTGAGCTGCTCCAACACTCTCTACCGAA
GATCAACCGGAGCGCTTCCGAGCCATCCTTGCATCGGGCAGCCCACACTGAGGATATCAAT
GCTTGCACGCTGACCACGTCCCCGAGGCTGCCTGTCTTCTGCGGCCGCTAGTCGACGGTAC
CGCGGGCCCGGGATAAGTCAACTAACTTAAGCTAGCAACGGTTTCCCTCTAGCGGGATCAA
TTcccccccccctaacgttactggccgaagccgcttggaataaggccggtgtgcgtttgt
ctatatgttattttccaccatattgccgtcttttggcaatgtgagggcccggaaacctggc
cctgtcttcttgacgagcattcctaggggtctttcccctctcgccaaaggaatgcaaggtc
tgttgaatgtcgtgaaggaagcagttcctctggaagcttcttgaagacaaacaacgtctgt
agcgaccctttgcaggcagcggaaccccccacctggcgacaggtgcctctgcggccaaaag
ccacgtgtataagatacacctgcaaaggcggcacaaccccagtgccacgttgtgagttgga
tagttgtggaaagagtcaaatggctctcctcaagcgtattcaacaaggggctgaaggatgc
ccagaaggtaccccattgtatgggatctgatctggggcctcggtgcacatgctttacatgt
gtttagtcgaggttaaaaaacgtctaggccccccgaaccacggggacgtggttttcctttg
aaaaacacgatAATACCATGACCGAGTACAAGCCCACGGTGCGCCTCGCCACCCGCGACGA
CGTCCCCAGGGCCGTACGCACCCTCGCCGCCGCGTTCGCCGACTACCCCGCCACGCGCCAC
ACCGTCGATCCGGACCGCCACATCGAGCGGGTCACCGAGCTGCAAGAACTCTTCCTCACGC
GCGTCGGGCTCGACATCGGCAAGGTGTGGGTCGCGGACGACGGCGCCGCGGTGGCGGTCTG
GACCACGCCGGAGAGCGTCGAAGCGGGGGCGGTGTTCGCCGAGATCGGCCCGCGCATGGCC
GAGTTGAGCGGTTCCCGGCTGGCCGCGCAGCAACAGATGGAAGGCCTCCTGGCGCCGCACC
GGCCCAAGGAGCCCGCGTGGTTCCTGGCCACCGTCGGCGTCTCGCCCGACCACCAGGGCAA
GGGTCTGGGCAGCGCCGTCGTGCTCCCCGGAGTGGAGGCGGCCGAGCGCGCCGGGGTGCCC
GCCTTCCTGGAGACCTCCGCGCCCCGCAACCTCCCCTTCTACGAGCGGCTCGGCTTCACCG
TCACCGCCGACGTCGAGTGCCCGAAGGACCGCGCGACCTGGTGCATGACCCGCAAGCCCGG
TGCCTGA

**j**

pPBbsr-ERK-KTR-mCherry

```
>Amino acid sequence
MKGRKPRDLELPLSPSLLGGQGPERTPGSGTSSGLQAPGPALSPSKRSGLEDPATPSKKPR
TPSVSSRLERLTLQSSFQFPSGGRMVSKGEEDNMAIIKEFMRFKVHMEGSVNGHEFEIEGE
GEGRPYEGTQTAKLKVTKGGPLPFAWDILSPQFMYGSKAYVKHPADIPDYLKLSFPEGFKW
ERVMNFEDGGVVTVTQDSSLQDGEFIYKVKLRGTNFPSDGPVMQKKTMGWEASSERMYPED
GALKGEIKQRLKLKDGGHYDAEVKTTYKAKKPVQLPGAYNVNIKLDITSHNEDYTIVEQYE
RAEGRHSTGGMDELYK*-[EMCV  IRES]-MVMKTFNISQQDLELVEVATEKITMLYEDNK
HHVGAAIRTKTGEIISAVHIEAYIGRVTVCAEAIAIGSAVSNGQKDFDTIVAVRHPYSDEV
DRSIRVVSPCGMCRELISDYAPDCFVLIEMNGKLVKTTIEELIPLKYTRN*

>DNA sequence
ATGAAAGGCCGCAAGCCTCGCGATCTGGAGTTACCTCTCAGCCCTAGCCTCCTGGGGGGAC
AGGGCCCTGAGCGCACACCCGGGTCCGGCACCTCTAGTGGTTTGCAGGCTCCGGGACCTGC
CCTGTCACCCTCCAAGCGCAGCGGCCTGGAGGATCCTGCCACACCCAGCAAAAAACCAAGG
ACCCCCTCAGTCAGCTCGCGGTTAGAGCGACTGACGCTCCAGTCTTCTTTCCAATTCCCGT
CCGGCGGCCGCATGGTGAGCAAGGGCGAGGAGGATAACATGGCCATCATCAAGGAGTTCAT
GCGCTTCAAGGTGCACATGGAGGGCTCCGTGAACGGCCACGAGTTCGAGATCGAGGGCGAG
GGCGAGGGCCGCCCCTACGAGGGCACCCAGACCGCCAAGCTGAAGGTGACCAAGGGTGGCC
CCCTGCCCTTCGCCTGGGACATCCTGTCCCCTCAGTTCATGTACGGCTCCAAGGCCTACGT
GAAGCACCCCGCCGACATCCCCGACTACTTGAAGCTGTCCTTCCCCGAGGGCTTCAAGTGG
GAGCGCGTGATGAACTTCGAGGACGGCGGCGTGGTGACCGTGACCCAGGACTCCTCCCTGC
AGGACGGCGAGTTCATCTACAAGGTGAAGCTGCGCGGCACCAACTTCCCCTCCGACGGCCC
CGTAATGCAGAAGAAGACCATGGGCTGGGAGGCCTCCTCCGAGCGGATGTACCCCGAGGAC
GGCGCCCTGAAGGGCGAGATCAAGCAGAGGCTGAAGCTGAAGGACGGCGGCCACTACGACG
CTGAGGTCAAGACCACCTACAAGGCCAAGAAGCCCGTGCAGCTGCCCGGCGCCTACAACGT
CAACATCAAGTTGGACATCACCTCCCACAACGAGGACTACACCATCGTGGAACAGTACGAA
CGCGCCGAGGGCCGCCACTCCACCGGCGGCATGGACGAGCTGTACAAGTAATCCGGACTCA
GATCTCGACAAGGTAGTGGTGCTGGCTCTGGTGCTGGTAGTGGCGCTGGTTCCGGTGCTGG
CTCTGGCGCGCCTCGAAGCCTCGAGATGGAGCACATACAGGGAGCTTGGAAGACGATCAGC
AATGGTTTTGGATTCAAAGATGCCGTGTTTGATGGCTCCAGCTGCATCTCTCCTACAATAG
TTCAGCAGTTTGGCTATCAGCGCCGGGCATCAGATGATGGCAAACTCACAGATCCTTCTAA
GACAAGCAACACTATCCGTGTTTTCTTGCCGAACAAGCAAAGAACAGTGGTCAATGTGCGA
AATGGAATGAGCTTGCATGACTGCCTTATGAAAGCACTCAAGGTGAGGGGCCTGCAACCAG
AGTGCTGTGCAGTGTTCAGACTTCTCCACGAACACAAAGGTAAAAAAGCACGCTTAGATTG
GAATACTGATGCTGCGTCTTTGATTGGAGAAGAACTTCAAGTAGATTTCCTGGATCATGTT
CCCCTCACAACACACAACTTTGCTCGGAAGACGTTCCTGAAGCTTGCCTTCTGTGACATCT
GTCAGAAATTCCTGCTCAATGGATTTCGATGTCAGACTTGTGGCTACAAATTTCATGAGCA
CTGTAGCACCAAAGTACCTACTATGTGTGTGGACTGGAGTAACATCAGACAACTCTTATTG
TTTCCAAATTCCACTATTGGTGATAGTGGAGTCCCAGCACTACCTTCTTTGACTATGCGTC
GTATGCGAGAGTCTGTTTCCAGGATGCCTGTTAGTTCTCAGCACAGATATTCTACACCTCA
CGCCTTCACCTTTAACACCTCCAGTCCCTCATCTGAAGGTTCCCTCTCCCAGAGGCAGAGG
TCGACATCCACACCTAATGTCCACATGGTCAGCACCACGCTGCCTGTGGACAGCAGGATGA
TTGAGGATGCAATTCGAAGTCACAGCGAATCAGCCTCACCTTCAGCCCTGTCCAGTAGCCC
CAACAATCTGAGCCCAACAGGCTGGTCACAGCCGAAAACCCCCGTGCCAGCACAAAGAGAG
CGGGCACCAGTATCTGGGACCCAGGAGAAAAACAAAATTAGGCCTCGTGGACAGAGAGATT
CAAGCTATTATTGGGAAATAGAAGCCAGTGAAGTGATGCTGTCCACTCGGATTGGGTCAGG
CTCTTTTGGAACTGTTTATAAGGGTAAATGGCACGGAGATGTTGCAGTAAAGATCCTAAAG
GTTGTCGACCCAACCCCAGAGCAATTCCAGGCCTTCAGGAATGAGGTGGCTGTTCTGCGCA
AAACACGGCATGTGAACATTCTGCTTTTCATGGGGTACATGACAAAGGACAACCTGGCAAT
TGTGACCCAGTGGTGCGAGGGCAGCAGCCTCTACAAACACCTGCATGTCCAGGAGACCAAG
```

TTTCAGATGTTCCAGCTAATTGACATTGCCCGGCAGACGGCTCAGGGAATGGACTATTTGC
ATGCAAAGAACATCATCCATAGAGACATGAAATCCAACAATATATTTCTCCATGAAGGCTT
AACAGTGAAAATTGGAGATTTTGGTTTGGCAACAGTAAAGTCACGCTGGAGTGGTTCTCAG
CAGGTTGAACAACCTACTGGCTCTGTCCTCTGGATGGCCCCAGAGGTGATCCGAATGCAGG
ATAACAACCCATTCAGTTTCCAGTCGGATGTCTACTCCTATGGCATCGTATTGTATGAACT
GATGACGGGGGAGCTTCCTTATTCTCACATCAACAACCGAGATCAGATCATCTTCATGGTG
GGCCGAGGATATGCCTCCCCAGATCTTAGTAAGCTATATAAGAACTGCCCCAAAGCAATGA
AGAGGCTGGTAGCTGACTGTGTGAAGAAAGTAAAGGAAGAGAGGCCTCTTTTTCCCCAGAT
CCTGTCTTCCATTGAGCTGCTCCAACACTCTCTACCGAAGATCAACCGGAGCGCTTCCGAG
CCATCCTTGCATCGGGCAGCCCACACTGAGGATATCAATGCTTGCACGCTGACCACGTCCC
CGAGGCTGCCTGTCTTCGTCGACGGGCCGCGGTAACAATTGTTAACTAACTTAAGCTAGCA
ACGGTTTCCCTCTAGCGGGATCAATTCCGcccccccccctaacgttactggccgaagccg
cttggaataaggccggtgtgcgtttgtctatatgttattttccaccatattgccgtctttt
ggcaatgtgagggcccggaaacctggccctgtcttcttgacgagcattcctaggggtcttt
cccctctcgccaaaggaatgcaaggtctgttgaatgtcgtgaaggaagcagttcctctgga
agcttcttgaagacaaacaacgtctgtagcgaccctttgcaggcagcggaaccccccacct
ggcgacaggtgcctctgcggccaaaagccacgtgtataagatacacctgcaaaggcggcac
aaccccagtgccacgttgtgagttggatagttgtggaaagagtcaaatggctctcctcaag
cgtattcaacaaggggctgaaggatgcccagaaggtaccccattgtatgggatctgatctg
gggcctcggtgcacatgctttacatgtgtttagtcgaggttaaaaaacgtctaggccccc
gaaccacggggacgtggttttcctttgaaaaacacgatAATACCATGGTCATGAAAACATT
TAACATTTCTCAACAAGATCTAGAATTAGTAGAAGTAGCGACAGAGAAGATTACAATGCTT
TATGAGGATAATAAACATCATGTGGGAGCGGCAATTCGTACGAAAACAGGAGAAATCATTT
CGGCAGTACATATTGAAGCGTATATAGGACGAGTAACTGTTTGTGCAGAAGCCATTGCGAT
TGGTAGTGCAGTTTCGAATGGACAAAAGGATTTTGACACGATTGTAGCTGTTAGACACCCT
TATTCTGACGAAGTAGATAGAAGTATTCGAGTGGTAAGTCCTTGTGGTATGTGTAGGGAGT
TGATTTCAGACTATGCACCAGATTGTTTTGTGTTAATAGAAATGAATGGCAAGTTAGTCAA
AACTACGATTGAAGAACTCATTCCACTCAAATATACCCGAAATTAA

**k**

pCMV-K6-SNAP$_i$-EGFP-iSH2

>Amino acid sequence
MKKKKKKGSGADKDCEMKRTTLDSPLGKLELSGCEQGLHRIIFLGKGTSAADAVEVPAPAA
VLGGPEPLMQATAWLNAYFHQPEAIEEFPVPALHHPVFQQESFTRQVLWKLLKVVKFGEVI
SYSHLAALAGNPAATAAVKTALSGNPVPILIPCHRVVQGDLDVGGYEGGLAVKEWLLAHEA
AASDPPVATMVSKGEELFTGVVPILVELDGDVNGHKFSVSGEGEGDATYGKLTLKFICTTG
KLPVPWPTLVTTLTYGVQCFSRYPDHMKQHDFFKSAMPEGYVQERTIFFKDDGNYKTRAEV
KFEGDTLVNRIELKGIDFKEDGNILGHKLEYNYNSHNVYIMADKQKNGIKVNFKIRHNIED
GSVQLADHYQQNTPIGDGPVLLPDNHYLSTQSALSKDPNEKRDHMVLLEFVTAAGITLGMD
ELYKSGLRSRQGSGAGSGAGSGAGSGAGSGAPRASKYQQDQVVKEDSVEAVGAQLKVYHQQ
YQDKSREYDQLYEEYTRTSQELQMKRTAIEAFNETIKIFEEQGQTQEKCSKEYLERFRREG
NEKEMQRILLNSERLKSRIAEIHESRTKLEQDLRAQASDNREIDKRMNSLKPDLMQLRKIR
DQYLVWLTQKGARQRKINEWLGIKNETEDQYSLMEDEDALPHHEERT*

>DNA sequence
ATGAAAAAAAAGAAAAAGAAAGGCAGCGGTGCTGACAAAGACTGCGAAATGAAGCGCACCA
CCCTGGATAGCCCTCTGGGCAAGCTGGAACTGTCTGGGTGCGAACAGGGCCTGCACCGTAT
CATCTTCCTGGGCAAAGGAACATCTGCCGCCGACGCCGTGGAAGTGCCTGCCCCAGCCGCC
GTGCTGGGCGGACCAGAGCCACTGATGCAGGCCACCGCCTGGCTCAACGCCTACTTTCACC
AGCCTGAGGCCATCGAGGAGTTCCCTGTGCCAGCCCTGCACCACCCAGTGTTCCAGCAGGA
GAGCTTTACCCGCCAGGTGCTGTGGAAACTGCTGAAAGTGGTGAAGTTCGGAGAGGTCATC
AGCTACAGCCACCTGGCCGCCCTGGCCGGCAATCCCGCCGCCACCGCCGCCGTGAAAACCG
CCCTGAGCGGAAATCCCGTGCCCATTCTGATCCCCTGCCACCGGGTGGTGCAGGGCGACCT
GGACGTGGGGGGCTACGAGGGCGGGCTCGCCGTGAAAGAGTGGCTGCTGGCCCACGAGGCG
GCCGCTTCGGACCCACCGGTCGCCACCATGGTGAGCAAGGGCGAGGAGCTGTTCACCGGGG
TGGTGCCCATCCTGGTCGAGCTGGACGGCGACGTAAACGGCCACAAGTTCAGCGTGTCCGG
CGAGGGCGAGGGCGATGCCACCTACGGCAAGCTGACCCTGAAGTTCATCTGCACCACCGGC
AAGCTGCCCGTGCCCTGGCCCACCCTCGTGACCACCCTGACCTACGGCGTGCAGTGCTTCA
GCCGCTACCCCGACCACATGAAGCAGCACGACTTCTTCAAGTCCGCCATGCCCGAAGGCTA
CGTCCAGGAGCGCACCATCTTCTTCAAGGACGACGGCAACTACAAGACCCGCGCCGAGGTG
AAGTTCGAGGGCGACACCCTGGTGAACCGCATCGAGCTGAAGGGCATCGACTTCAAGGAGG
ACGGCAACATCCTGGGGCACAAGCTGGAGTACAACTACAACAGCCACAACGTCTATATCAT
GGCCGACAAGCAGAAGAACGGCATCAAGGTGAACTTCAAGATCCGCCACAACATCGAGGAC
GGCAGCGTGCAGCTCGCCGACCACTACCAGCAGAACACCCCCATCGGCGACGGCCCCGTGC
TGCTGCCCGACAACCACTACCTGAGCACCCAGTCCGCCCTGAGCAAAGACCCCAACGAGAA
GCGCGATCACATGGTCCTGCTGGAGTTCGTGACCGCCGCCGGGATCACTCTCGGCATGGAC
GAGCTGTACAAGTCCGGACTCAGATCTCGACAAGGTAGTGGTGCTGGCTCTGGTGCTGGTA
GTGGCGCTGGTTCCGGTGCTGGCTCTGGCGCGCCTCGAGCATCCAAGTACCAACAAGACCA
GGTGGTGAAGGAGGACAGCGTAGAGGCTGTGGGCGCCCAGCTCAAGGTCTACCACCAGCAG
TACCAGGACAAGAGCCGCGAATATGACCAGCTGTATGAAGAATACACACGGACCTCCCAGG
AGCTGCAGATGAAGCGCACAGCCATAGAGGCCTTCAACGAGACCATCAAGATCTTCGAAGA
GCAGGGCCAGACACAGGAGAAGTGCAGCAAGGAGTATTTGGAGCGCTTCCGGCGAGAGGGA
AATGAGAAGGAGATGCAGAGGATCCTGCTGAACTCCGAGCGACTCAAGTCTCGCATCGCGG
AGATACACGAAAGCCGCACGAAGTTGGAGCAGGATCTGCGGGCGCAGGCCTCCGACAACCG
TGAGATCGACAAGCGCATGAACAGCCTCAAACCTGACCTCATGCAGCTGCGCAAGATCAGG
GACCAGTACCTCGTGTGGCTCACCCAGAAAGGTGCCCGACAGAGGAAGATCAACGAATGGC
TGGGAATCAAGAACGAGACTGAGGACCAGTATTCACTGATGGAGGATGAGGACGCCCTCCC
CCACCACGAGGAGCGCACGTGA

**I**

pCMV-mCherry-$PH_{Akt}$

```
>Amino acid sequence
MVSKGEEDNMAIIKEFMRFKVHMEGSVNGHEFEIEGEGEGRPYEGTQTAKLKVTKGGPLPF
AWDILSPQFMYGSKAYVKHPADIPDYLKLSFPEGFKWERVMNFEDGGVVTVTQDSSLQDGE
FIYKVKLRGTNFPSDGPVMQKKTMGWEASSERMYPEDGALKGEIKQRLKLKDGGHYDAEVK
TTYKAKKPVQLPGAYNVNIKLDITSHNEDYTIVEQYERAEGRHSTGGMDELYK  SGLRSRA
QASNSAVDGTAGPGSMSDVAIVKEGWLHKRGEYIKTWRPRYFLLKNDGTFIGYKERPQDVD
QREAPLNNFSVAQCQLMKTERPRPNTFIIRCLQWTTVIERTFHVETPEEREEWTTAIQTVA
DGLKKQEEEEMDFRSGSPSDNSGAEEMEVSLAKPKHRVTMN*

>DNA sequence
ATGGTGAGCAAGGGCGAGGAGGATAACATGGCCATCATCAAGGAGTTCATGCGCTTCAAGG
TGCACATGGAGGGCTCCGTGAACGGCCACGAGTTCGAGATCGAGGGCGAGGGCGAGGGCCG
CCCCTACGAGGGCACCCAGACCGCCAAGCTGAAGGTGACCAAGGGTGGCCCCCTGCCCTTC
GCCTGGGACATCCTGTCCCCTCAGTTCATGTACGGCTCCAAGGCCTACGTGAAGCACCCCG
CCGACATCCCCGACTACTTGAAGCTGTCCTTCCCCGAGGGCTTCAAGTGGGAGCGCGTGAT
GAACTTCGAGGACGGCGGCGTGGTGACCGTGACCCAGGACTCCTCCCTGCAGGACGGCGAG
TTCATCTACAAGGTGAAGCTGCGCGGCACCAACTTCCCCTCCGACGGCCCCGTAATGCAGA
AGAAGACCATGGGCTGGGAGGCCTCCTCCGAGCGGATGTACCCCGAGGACGGCGCCCTGAA
GGGCGAGATCAAGCAGAGGCTGAAGCTGAAGGACGGCGGCCACTACGACGCTGAGGTCAAG
ACCACCTACAAGGCCAAGAAGCCCGTGCAGCTGCCCGGCGCCTACAACGTCAACATCAAGT
TGGACATCACCTCCCACAACGAGGACTACACCATCGTGGAACAGTACGAACGCGCCGAGGG
CCGCCACTCCACCGGCGGCATGGACGAGCTGTACAAGTCCGGACTCAGATCTCGAGCTCAA
GCTTCGAATTCTGCAGTCGACGGTACCGCGGGCCCGGGATCCATGAGCGACGTGGCTATTG
TGAAGGAGGGTTGGCTGCACAAACGAGGGGAGTACATCAAGACCTGGCGGCCACGCTACTT
CCTCCTCAAGAATGATGGCACCTTCATTGGCTACAAGGAGCGGCCGCAGGATGTGGACCAA
CGTGAGGCTCCCCTCAACAACTTCTCTGTGGCGCAGTGCCAGCTGATGAAGACGGAGCGGC
CCCGGCCCAACACCTTCATCATCCGCTGCCTGCAGTGGACCACTGTCATCGAACGCACCTT
CCATGTGGAGACTCCTGAGGAGCGGGAGGAGTGGACAACCGCCATCCAGACTGTGGCTGAC
GGCCTCAAGAAGCAGGAGGAGGAGGAGATGGACTTCCGGTCGGGCTCACCCAGTGACAACT
CAGGGGCTGAAGAGATGGAGGTGTCCCTGGCCAAGCCCAAGCACCGCGTGACCATGAACTA
A
```

**m**

pCMV-K6-SNAP$_i$-EGFP-Sos

>Amino acid sequence
MKKKKKKGSGADKDCEMKRTTLDSPLGKLELSGCEQGLHRIIFLGKGTSAADAVEVPAPAA
VLGGPEPLMQATAWLNAYFHQPEAIEEFPVPALHHPVFQQESFTRQVLWKLLKVVKFGEVI
SYSHLAALAGNPAATAAVKTALSGNPVPILIPCHRVVQGDLDVGGYEGGLAVKEWLLAHEA
AASDPPVATMVSKGEELFTGVVPILVELDGDVNGHKFSVSGEGEGDATYGKLTLKFICTTG
KLPVPWPTLVTTLTYGVQCFSRYPDHMKQHDFFKSAMPEGYVQERTIFFKDDGNYKTRAEV
KFEGDTLVNRIELKGIDFKEDGNILGHKLEYNYNSHNVYIMADKQKNGIKVNFKIRHNIED
GSVQLADHYQQNTPIGDGPVLLPDNHYLSTQSALSKDPNEKRDHMVLLEFVTAAGITLGMD
ELYKSGLRSRQGSGAGSGAGSGAGSGAGSGAPRAQASQAQQLPYEFFSEENAPKWRGLLVP
ALKKVQGQVHPTLESNDDALQYVEELILQLLNMLCQAQPRSASDVEERVQKSFPHPIDKWA
IADAQSAIEKRKRRNPLSLPVEKIHPLLKEVLGYKIDHQVSVYIVAVLEYISADILKLVGN
YVRNIRHYEITKQDIKVAMCADKVLMDMFHQDVEDINILSLTDEEPSTSGEQTYYDLVKAF
MAEIRQYIRELNLIIKVFREPFVSNSKLFSANDVENIFSRIVDIHELSVKLLGHIEDTVEM
TDEGSPHPLVGSCFEDLAEELAFDPYESYARDILRPGFHDRFLSQLSKPGAALYLQSIGEG
FKEAVQYVLPRLLLAPVYHCLHYFELLKQLEEKSEDQEDKECLKQAITALLNVQSGMEKIC
SKSLAKRRLSESACRFYSQQMKGKQLAIKKMNEIQKNIDGWEGKDIGQCCNEFIMEGTLTR
VGAKHERHIFLFDGLMICCKSNHGQPRLPGASNAEYRLKEKFFMRKVQINDKDDTNEYKHA
FEIILKDENSVIFSAKSAEEKNNWMAALISLQYRSTLERMLDVTMLQEEKEEQMRLPSADV
YRFAEPDSEENIIFEENMQPKAGIPIIKAGTVIKLIERLTYHMYADPNFVRTFLTTYRSFC
KPQELLSLIIERFEIPEPEPTEADRIAIENGDQPLSAELKRFRKEYIQPVQLRVLNVCRHW
VEHHFYDFERDAYLLQRMEEFIGTVRGKAMKKWVESITKIIQRKKIARDNGPGHNITFQSS
PPTVEWHISRPGHIETFDLLTLHPIEIARQLTLLESDLYRAVQPSELVGSVWTKEDKEINS
PNLLKMIRHTTNLTLWFEKCIVETENLEERVAVVSRIIEILQVFQELNNFNGVLEVVSAMN
SSPVYRLDHTFEQIPSRQKKILEEAHELSEDHYKKYLAKLRSINPPCVPFFGIYLTNILKT
EEGNPEVLKRHGKELINFSKRRKVAEITGEIQQYQNQPYCLRVESDIKRFFENLNPMGNSM
EKEFTDYLFNKSLEIEPRNPKPLPRFPKKYSYPLKSPGVRPSNPRPGTMRHPTPLQQEPRK
ISYSRIPESETESTASAPNSPRTPLTPPPASGASSTTDVCSVFDSDHSSPFHSSNDTVFIQ
VTLPHGPRSASVSSISLTKGTDEVPVPPPVPPRRRPESAPAESSPSKIMSKHLDSPPAIPP
RQPTSKAYSPRYSISDRTSISDPPESPPLLPPREPVRTPDVFSSSPLHLQPPPLGKKSDHG
NAFFPNSPSPFTPPPPQTPSPHGTRRHLPSPPLTQEVDLHSIAGPPVPPRQSTSQHIPKLP
PKTYKREHTHPSMHRDGPPLLENAHSSPRARDPPDLDN*

>DNA sequence
ATGAAAAAAAAGAAAAAGAAAGGCAGCGGTGCTGACAAAGACTGCGAAATGAAGCGCACCA
CCCTGGATAGCCCTCTGGGCAAGCTGGAACTGTCTGGGTGCGAACAGGGCCTGCACCGTAT
CATCTTCCTGGGCAAAGGAACATCTGCCGCCGACGCCGTGGAAGTGCCTGCCCCAGCCGCC
GTGCTGGGCGGACCAGAGCCACTGATGCAGGCCACCGCCTGGCTCAACGCCTACTTTCACC
AGCCTGAGGCCATCGAGGAGTTCCCTGTGCCAGCCCTGCACCACCCAGTGTTCCAGCAGGA
GAGCTTTACCCGCCAGGTGCTGTGGAAACTGCTGAAAGTGGTGAAGTTCGGAGAGGTCATC
AGCTACAGCCACCTGGCCGCCCTGGCCGGCAATCCCGCCGCCACCGCCGCCGTGAAAACCG
CCCTGAGCGGAAATCCCGTGCCCATTCTGATCCCCTGCCACCGGGTGGTGCAGGGCGACCT
GGACGTGGGGGGCTACGAGGGCGGGCTCGCCGTGAAAGAGTGGCTGCTGGCCCACGAGGCG
GCCGCTTCGGACCCACCGGTCGCCACCATGGTGAGCAAGGGCGAGGAGCTGTTCACCGGGG
TGGTGCCCATCCTGGTCGAGCTGGACGGCGACGTAAACGGCCACAAGTTCAGCGTGTCCGG
CGAGGGCGAGGGCGATGCCACCTACGGCAAGCTGACCCTGAAGTTCATCTGCACCACCGGC
AAGCTGCCCGTGCCCTGGCCCACCCTCGTGACCACCCTGACCTACGGCGTGCAGTGCTTCA
GCCGCTACCCCGACCACATGAAGCAGCACGACTTCTTCAAGTCCGCCATGCCCGAAGGCTA
CGTCCAGGAGCGCACCATCTTCTTCAAGGACGACGGCAACTACAAGACCCGCGCCGAGGTG
AAGTTCGAGGGCGACACCCTGGTGAACCGCATCGAGCTGAAGGGCATCGACTTCAAGGAGG
ACGGCAACATCCTGGGGCACAAGCTGGAGTACAACTACAACAGCCACAACGTCTATATCAT

GGCCGACAAGCAGAAGAACGGCATCAAGGTGAACTTCAAGATCCGCCACAACATCGAGGAC
GGCAGCGTGCAGCTCGCCGACCACTACCAGCAGAACACCCCCATCGGCGACGGCCCCGTGC
TGCTGCCCGACAACCACTACCTGAGCACCCAGTCCGCCCTGAGCAAAGACCCCAACGAGAA
GCGCGATCACATGGTCCTGCTGGAGTTCGTGACCGCCGCCGGGATCACTCTCGGCATGGAC
GAGCTGTACAAGTCCGGACTCAGATCTCGACAAGGTAGTGGTGCTGGCTCTGGTGCTGGTA
GTGGCGCTGGTTCCGGTGCTGGCTCTGGCGCGCCTCGAGCTCAAGCTTCGCAGGCGCAGCA
GCTGCCCTACGAGTTTTTCAGCGAAGAGAACGCGCCCAAGTGGCGGGGACTACTGGTGCCT
GCGCTGAAAAAGGTCCAGGGGCAAGTTCATCCTACTCTCGAGTCTAATGATGATGCTCTTC
AGTATGTTGAAGAATTAATTTTGCAATTATTAAATATGCTATGCCAAGCTCAGCCCCGAAG
TGCTTCAGATGTAGAGGAACGTGTTCAAAAAAGTTTCCCTCATCCAATTGATAAATGGGCA
ATAGCTGATGCCCAATCAGCTATTGAAAAGAGGAAGCGAAGAAACCCTTTATCTCTCCCAG
TAGAAAAAATTCATCCTTTATTAAAGGAGGTCCTAGGTTATAAAATTGACCACCAGGTTTC
TGTTTACATAGTAGCAGTCTTAGAATACATTTCTGCAGACATTTTAAAGCTGGTTGGGAAT
TATGTAAGAAATATACGGCATTATGAAATTACAAAACAAGATATTAAAGTGGCAATGTGTG
CTGACAAGGTATTGATGGATATGTTTCATCAAGATGTAGAAGATATTAATATATTATCTTT
AACTGACGAAGAGCCTTCCACCTCAGGAGAACAAACTTACTATGATTTGGTAAAAGCATTT
ATGGCAGAAATTCGACAATATATAAGGGAACTAAATCTAATTATAAAAGTTTTTAGAGAGC
CCTTTGTCTCCAATTCAAAATTGTTTTCAGCTAATGATGTAGAAAATATATTTAGTCGCAT
AGTAGATATACATGAACTTAGTGTAAAGTTACTGGGCCATATAGAAGATACAGTAGAAATG
ACAGATGAAGGCAGTCCCCATCCACTAGTAGGAAGCTGCTTTGAAGACTTAGCAGAGGAAC
TGGCATTTGATCCATATGAATCGTATGCTCGAGATATTTTGCGACCTGGTTTTCATGATCG
TTTCCTTAGTCAGTTATCAAAGCCTGGGGCAGCACTTTATTTGCAGTCAATAGGCGAAGGT
TTCAAAGAAGCTGTTCAATATGTTTTACCCAGGCTGCTTCTGGCCCCTGTTTACCACTGTC
TCCATTACTTTGAACTTTTGAAGCAGTTAGAAGAAAAAAGTGAAGATCAAGAAGACAAGGA
ATGTTTAAAACAAGCAATAACAGCTTTGCTTAATGTTCAGAGTGGTATGGAAAAAATATGT
TCTAAAAGTCTTGCAAAACGAAGACTGAGTGAATCTGCATGTCGGTTTTATAGTCAGCAAA
TGAAGGGGAAACAACTAGCAATCAAGAAGATGAACGAGATTCAGAAGAATATTGATGGTTG
GGAGGGAAAAGACATTGGACAGTGTTGTAATGAATTTATAATGGAAGGAACTCTTACACGT
GTAGGAGCCAAACATGAGAGACACATATTTCTCTTTGATGGCTTAATGATTTGCTGTAAAT
CAAATCATGGGCAGCCAAGACTTCCTGGTGCTAGCAATGCAGAATATCGTCTTAAAGAAAA
GTTTTTTATGCGAAAGGTACAAATTAATGATAAAGATGACACCAATGAATACAAGCATGCT
TTTGAAATAATTTTAAAAGATGAAAATAGTGTTATATTTTCTGCCAAGTCAGCTGAAGAGA
AAAACAATTGGATGGCAGCATTGATATCTTTACAGTACCGGAGTACACTGGAAAGGATGCT
TGATGTAACAATGCTACAGGAAGAGAAAGAGGAGCAGATGAGGCTGCCTAGTGCTGATGTT
TATAGATTTGCAGAGCCTGACTCTGAAGAGAATATTATATTTGAAGAGAACATGCAGCCCA
AGGCTGGAATTCCAATTATCAAAGCAGGAACTGTTATTAAACTTATAGAGAGGCTTACGTA
CCATATGTACGCAGATCCCAATTTTGTTCGGACATTTCTTACAACATACAGATCCTTTTGC
AAACCTCAAGAACTACTGAGTCTTATAATAGAAAGGTTTGAAATTCCAGAGCCTGAGCCAA
CAGAAGCTGATCGCATAGCTATAGAGAATGGAGATCAACCCTTGAGTGCAGAACTGAAAAG
ATTTAGAAAAGAATATATACAGCCTGTGCAACTGCGAGTATTAAATGTATGTCGGCACTGG
GTAGAGCACCACTTCTATGATTTTGAAAGAGATGCATATCTTTTGCAACGAATGGAAGAAT
TTATTGGAACAGTAAGAGGTAAAGCAATGAAAAAATGGGTTGAATCCATCACTAAAATAAT
CCAAAGGAAAAAAATTGCAAGAGACAATGGACCAGGTCATAATATTACATTTCAGAGTTCA
CCTCCCACAGTTGAGTGGCATATAAGCAGACCTGGGCACATAGAGACTTTTGACCTGCTCA
CCTTACACCCAATAGAAATTGCTCGACAACTCACTTTACTTGAATCAGATCTATACCGAGC
TGTACAGCCATCAGAATTAGTTGGAAGTGTGTGGACAAAAGAAGACAAAGAAATTAACTCT
CCTAATCTTCTGAAAATGATTCGACATACCACCAACCTCACTCTGTGGTTTGAGAAATGTA
TTGTAGAAACTGAAAATTTAGAAGAAAGAGTAGCTGTGGTGAGTCGAATTATTGAGATTCT
ACAAGTCTTTCAAGAGTTGAACAACTTTAATGGTGTCCTTGAGGTTGTCAGTGCTATGAAT
TCATCACCTGTTTACAGACTAGACCACACATTTGAGCAAATACCAAGTCGCCAGAAGAAAA
TTTTAGAAGAAGCTCATGAATTGAGTGAAGATCACTATAAGAAATATTTGGCAAAACTCAG
GTCTATTAATCCACCATGTGTGCCTTTCTTTGGAATTTATCTCACTAATATCTTGAAAACA
GAAGAAGGCAACCCTGAGGTCCTAAAAAGACATGGAAAAGAGCTTATAAACTTTAGCAAAA
GGAGGAAAGTAGCAGAAATAACAGGAGAGATCCAGCAGTACCAAAATCAGCCTTACTGTTT

ACGAGTAGAATCAGATATCAAAAGGTTCTTTGAAAACTTGAATCCGATGGGAAATAGCATG
GAGAAGGAATTTACAGATTATCTTTTCAACAAATCCCTAGAAATAGAACCACGAAACCCTA
AGCCTCTCCCAAGATTTCCAAAAAAATATAGCTATCCCCTAAAATCTCCTGGTGTTCGTCC
ATCAAACCCAAGACCAGGTACCATGAGGCATCCCACACCTCTGCAGCAGGAGCCAAGGAAA
ATTAGTTATAGTAGGATCCCTGAAAGTGAAACAGAAAGTACAGCATCTGCACCAAATTCTC
CAAGAACACCGTTAACACCTCCGCCTGCTTCTGGTGCTTCCAGTACCACAGATGTTTGCAG
TGTATTTGATTCCGATCATTCGAGCCCTTTTCACTCAAGCAATGATACCGTCTTTATCCAA
GTTACTCTGCCCCATGGCCCAAGATCTGCTTCTGTATCATCTATAAGTTTAACCAAAGGCA
CTGATGAAGTGCCTGTCCCTCCTCCTGTTCCTCCACGAAGACGACCAGAATCTGCCCCAGC
AGAATCTTCACCATCTAAGATTATGTCTAAGCATTTGGACAGTCCCCCAGCCATTCCTCCT
AGGCAACCCACATCAAAAGCCTATTCACCACGATATTCAATATCAGACCGGACCTCTATCT
CAGACCCTCCTGAAAGCCCTCCCTTATTACCACCACGAGAACCTGTGAGGACACCTGATGT
TTTCTCAAGCTCACCACTACATCTCCAACCTCCCCCTTTGGGCAAAAAAAGTGACCATGGC
AATGCCTTCTTCCCAAACAGCCCTTCCCCCTTTACACCACCTCCTCCTCAAACACCTTCTC
CTCACGGCACAAGAAGGCATCTGCCATCACCACCATTGACACAAGAAGTGGACCTTCATTC
CATTGCTGGGCCGCCTGTTCCTCCACGACAAAGCACTTCTCAACATATCCCTAAACTCCCT
CCAAAAACTTACAAAAGGGAGCACACACACCCATCCATGCACAGAGATGGACCACCACTGT
TGGAGAATGCCCATTCTTCCCCGCGGGCCCGGGATCCACCGGATCTAGATAACTGA

**n**

pPBbsr-MEK-P2A-mCherry-ERK

```
>Amino acid sequence
MDYKDDDDKARLEMPKKKPTPIQLNPNPEGTAVNGTPTAETNLEALQKKLEELELDEQQRK
RLEAFLTQKQKVGELKDDDFEKVSELGAGNGGVVFKVSHKPTSLIMARKLIHLEIKPAIRN
QIIRELQVLHECNSPYIVGFYGAFYSDGEISICMEHMDGGSLDQVLKKAGKIPEKILGKVS
IAVIKGLTYLREKHKIMHRDVKPSNILVNSRGEIKLCDFGVSGQLIDSMANSFVGTRSYMS
PERLQGTHYSVQSDIWSMGLSLVEMAIGRYPIPPPDAKELELIFGCSVERDPASSELAPRP
RPPGRPISSYGPDSRPPMAIFELLDYIVNEPPPKLPSGVFGAEFQDFVNKCLVKNPAERAD
LKQLMVHSFIKQSELEEVDFAGWLCSTMGLKQPSTPTHAAGVGGRGTSGSGATNFSLLKQA
GDVEENPGPQLIKGAMVSKGEEDNMAIIKEFMRFKVHMEGSVNGHEFEIEGEGEGRPYEGT
QTAKLKVTKGGPLPFAWDILSPQFMYGSKAYVKHPADIPDYLKLSFPEGFKWERVMNFEDG
GVVTVTQDSSLQDGEFIYKVKLRGTNFPSDGPVMQKKTMGWEASSERMYPEDGALKGEIKQ
RLKLKDGGHYDAEVKTTYKAKKPVQLPGAYNVNIKLDITSHNEDYTIVEQYDRAEGRHSTG
GMDELYLEMAAAGAASNPGGGPEMVRGQAFDVGPRYINLAYIGEGAYGMVCSAHDNVNKVR
VAIRKISPFEHQTYCQRTLREIKILLRFKHENIIGINDIIRAPTIEQMKDVYIVQDLMETD
LYKLLKTQHLSNDHICYFLYQILRGLKYIHSANVLHRDLKPSNLLLNTTCDLKICDFGLAR
VADPDHDHTGFLTEYVATRWYRAPEIMLNSKGYTKSIDIWSVGCILAEMLSNRPIFPGKHY
LDQLNHILGILGSPSQEDLNCIINLKARNYLLSLPHKNKVPWNRLFPNADPKALDLLDKML
TFNPHKRIEVEAALAHPYLEQYYDPSDEPVAEAPFKFEMELDDLPKETLKELIFEETARFQ
PGY*-[EMCV IRES]-MLYEDNKHHVGAAIRTKTGEIISAVHIEAYIGRVTVCAEAIAIG
SAVSNGQKDFDTIVAVRHPYSDEVDRSIRVVSPCGMCRELISDYAPDCFVLIEMNGKLVKT
TIEELIPLKYTRN*
```

```
>DNA sequence
ATGGACTACAAAGACGATGACGATAAAGCAAGGCTCGAGATGCCTAAAAAGAAGCCTACGC
CCATACAGCTGAATCCCAACCCCGAAGGGACTGCTGTGAACGGGACCCCTACAGCCGAGAC
AAACCTTGAAGCTCTGCAGAAAAAGTTGGAAGAGCTTGAGCTGGATGAGCAGCAGAGGAAG
CGTCTGGAGGCTTTTCTCACCCAGAAGCAGAAAGTTGGGGAACTGAAGGATGACGACTTTG
AAAAAGTTTCAGAGCTTGGAGCAGGCAACGGAGGAGTGGTGTTTAAGGTGTCCCACAAGCC
AACCAGCTTGATTATGGCCAGGAAGTTGATTCATCTGGAGATTAAGCCTGCAATCCGAAAC
CAGATTATCCGAGAGTTGCAGGTTCTGCATGAATGTAACTCCCCATACATTGTGGGGTTCT
ATGGGGCCTTCTACAGTGATGGAGAGATCAGCATTTGCATGGAACACATGGATGGAGGCTC
CCTTGATCAGGTTCTGAAGAAAGCTGGCAAAATCCCAGAAAAGATTTTGGGAAAAGTCAGC
ATTGCAGTGATAAAAGGTCTAACCTACCTGAGAGAAAAGCATAAGATAATGCACAGAGATG
TGAAACCTTCTAACATCCTGGTCAACTCTAGAGGAGAGATAAAACTCTGCGACTTTGGGGT
CAGCGGGCAACTCATAGACTCCATGGCAAATTCCTTTGTTGGGACAAGATCCTATATGTCA
CCGGAGCGACTACAGGGCACTCATTATTCTGTGCAATCAGACATCTGGAGCATGGGGCTGT
CGCTGGTGGAAATGGCCATTGGAAGGTATCCCATTCCACCCCCTGATGCCAAAGAGCTGGA
ACTTATCTTTGGGTGTTCTGTAGAAAGGGATCCAGCGTCTTCTGAACTGGCACCTCGCCCC
CGGCCACCCGGACGTCCAATAAGCTCATACGGTCCTGATAGTCGACCACCCATGGCTATTT
TTGAACTTCTGGATTATATCGTGAACGAGCCGCCTCCAAAATTGCCCAGTGGAGTATTTGG
AGCTGAGTTCCAGGACTTTGTGAATAAATGTCTTGTGAAGAATCCGGCAGAGAGAGCAGAC
CTTAAACAGCTAATGGTTCACAGCTTCATTAAGCAGTCAGAGTTGGAGGAAGTGGATTTTG
CTGGATGGCTCTGTTCCACTATGGGCCTTAAGCAGCCCAGTACCCCAACCCATGCCGCCGG
AGTGGGCGGCCGCGGCACTAGTGGAAGCGGAGCTACTAACTTCAGCCTGCTGAAGCAGGCT
GGAGACGTGGAGGAGAACCCTGGACCTCAATTAATTAAGGGCGCAATGGTGAGCAAGGGCG
AGGAGGATAACATGGCCATCATCAAGGAGTTCATGCGCTTCAAGGTGCACATGGAGGGCTC
CGTGAACGGCCACGAGTTCGAGATCGAGGGCGAGGGCGAGGGCCGCCCCTACGAGGGCACC
CAGACCGCCAAGCTGAAGGTGACCAAGGGTGGCCCCCTGCCCTTCGCCTGGGACATCCTGT
CCCCTCAGTTCATGTACGGCTCCAAGGCCTACGTGAAGCACCCCGCCGACATCCCCGACTA
```

CTTGAAGCTGTCCTTCCCCGAGGGCTTCAAGTGGGAGCGCGTGATGAACTTCGAGGACGGC
GGCGTGGTGACCGTGACCCAGGACTCCTCCCTGCAGGACGGCGAGTTCATCTACAAGGTGA
AGCTGCGCGGCACCAACTTCCCCTCCGACGGCCCCGTAATGCAGAAGAAGACCATGGGCTG
GGAGGCCTCCTCCGAGCGGATGTACCCCGAGGACGGCGCCCTGAAGGGCGAGATCAAGCAG
AGGCTGAAGCTGAAGGACGGCGGCCACTACGACGCTGAGGTCAAGACCACCTACAAGGCCA
AGAAGCCCGTGCAGCTGCCCGGCGCCTACAACGTCAACATCAAGTTGGACATCACCTCCCA
CAACGAGGACTACACCATCGTGGAACAGTACGACCGCGCCGAGGGCCGCCACTCCACCGGC
GGCATGGACGAGCTGTACCTCGAGATGGCAGCGGCAGGAGCTGCGTCTAACCCCGGCGGGG
GTCCGGAGATGGTGCGGGGCCAGGCGTTCGACGTAGGCCCTCGATACATCAATCTGGCTTA
TATCGGCGAGGGAGCGTACGGCATGGTGTGTTCTGCCCATGACAATGTTAACAAAGTTCGA
GTTGCTATCAGGAAAATCAGCCCATTTGAGCATCAGACATACTGCCAGCGAACATTGCGGG
AGATCAAAATCTTGCTACGTTTTAAACATGAAAACATCATTGGGATAAACGACATTATTCG
CGCTCCAACCATTGAGCAGATGAAAGATGTGTACATTGTGCAGGACCTCATGGAGACAGAC
CTCTATAAGCTCCTGAAGACTCAGCATCTTAGCAATGACCATATCTGCTATTTCTTGTACC
AGATTCTGAGAGGATTAAAGTACATCCATTCAGCCAATGTTCTACATCGTGATCTTAAGCC
TTCAAATTTGCTGCTTAACACTACCTGTGATCTCAAGATCTGTGATTTTGGATTGGCTCGT
GTTGCAGACCCAGATCATGATCACACTGGCTTTCTCACAGAATATGTAGCCACTCGCTGGT
ACAGAGCTCCTGAGATCATGCTGAATTCCAAGGGCTATACCAAATCAATTGACATCTGGTC
TGTTGGCTGCATTCTTGCTGAGATGCTTTCTAATAGACCCATATTTCCTGGGAAACATTAT
CTTGACCAGCTTAATCACATACTTGGTATTCTTGGATCTCCATCTCAAGAGGACCTAAACT
GTATAATCAATTTAAAAGCTAGGAATTACTTGCTTTCCCTTCCTCACAAAAATAAGGTGCC
ATGGAACAGACTTTTCCCCAATGCAGATCCCAAAGCTCTAGACTTACTGGACAAGATGCTG
ACTTTCAACCCCCATAAAAGAATTGAAGTAGAGGCAGCTTTGGCTCATCCTTATCTGGAGC
AGTATTATGACCCAAGTGATGAGCCTGTAGCTGAAGCTCCCTTTAAATTTGAAATGGAGCT
TGATGATTTGCCCAAGGAGACTCTTAAGGAGCTAATTTTTGAAGAAACCGCTAGATTCCAG
CCAGGGTACTAATCGCGCCTCTAGAGGATCCGTTAACTAACTTAAGCTAGCGTCGACGGGC
CGCGGTAACAATTGTTAACTAACTTAAGCTAGCAACGGTTTCCCTCTAGCGGGATCAATTC
CGcccccccccctaacgttactggccgaagccgcttggaataaggccggtgtgcgtttgt
ctatatgttattttccaccatattgccgtcttttggcaatgtgagggcccggaaacctggc
cctgtcttcttgacgagcattcctaggggtctttcccctctcgccaaaggaatgcaaggtc
tgttgaatgtcgtgaaggaagcagttcctctggaagcttcttgaagacaaacaacgtctgt
agcgaccctttgcaggcagcggaaccccccacctggcgacaggtgcctctgcggccaaaag
ccacgtgtataagatacacctgcaaaggcggcacaaccccagtgccacgttgtgagttgga
tagttgtggaaagagtcaaatggctctcctcaagcgtattcaacaaggggctgaaggatgc
ccagaaggtaccccattgtatgggatctgatctggggcctcggtgcacatgctttacatgt
gtttagtcgaggttaaaaaacgtctaggccccccgaaccacggggacgtggttttcctttg
aaaaacacgatAATACCATGGTCATGAAAACATTTAACATTTCTCAACAAGATCTAGAATT
AGTAGAAGTAGCGACAGAGAAGATTACAATGCTTTATGAGGATAATAAACATCATGTGGGA
GCGGCAATTCGTACGAAAACAGGAGAAATCATTTCGGCAGTACATATTGAAGCGTATATAG
GACGAGTAACTGTTTGTGCAGAAGCCATTGCGATTGGTAGTGCAGTTTCGAATGGACAAAA
GGATTTTGACACGATTGTAGCTGTTAGACACCCTTATTCTGACGAAGTAGATAGAAGTATT
CGAGTGGTAAGTCCTTGTGGTATGTGTAGGGAGTTGATTTCAGACTATGCACCAGATTGTT
TTGTGTTAATAGAAATGAATGGCAAGTTAGTCAAAACTACGATTGAAGAACTCATTCCACT
CAAATATACCCGAAATTAA

**Fig. S8.** DNA and amino acid sequences of constructs used in this study. (**a**) pPBpuro-K6-SNAP$_f$-Halo: orange, K6-tag; purple, SNAP$_f$; gray box, HaloTag; yellow marker, an internal ribosomal entry site from Encephalomyocarditis virus (EMCV IRES)[S11]; brown, puromycin *N*-acetyltransferase (pac). (**b**) pPBpuro-K6-SNAP$_i$-Halo: orange, K6-tag; purple, SNAP$_i$; gray box, HaloTag; yellow marker, EMCV IRES; brown, pac. (**c**) pPBpuro-K6-SNAP$_f$-ns-Halo: orange, K6-tag; purple, SNAP$_f$; gray box, HaloTag; yellow marker, EMCV IRES; brown, pac. (**d**) pPBpuro-K6-SNAP$_f$-EGFP: orange, K6-tag; purple, SNAP$_f$; green, EGFP; yellow marker,

EMCV IRES; brown, pac. (**e**) pPBpuro-K6-SNAP$_i$-EGFP: orange, K6-tag; purple, SNAP$_i$; green, EGFP; yellow marker, EMCV IRES; brown, pac. (**f**) pCMV-K6-SNAP$_f$-EGFP-Tiam1: orange, K6-tag; purple, SNAP$_f$; green, EGFP; gray box, a DH-PH domain (residues 1012–1591) from human Tiam1. (**g**) pCMV-K6-SNAP$_i$-EGFP-Tiam1: orange, K6-tag; purple, SNAP$_i$; green, EGFP; gray box, a DH-PH domain (residues 1012–1591) from human Tiam1. (**h**) pCMV-Lifeact-mCherry: gray box, Lifeact; red, mCherry. (**i**) pPBpuro-K6-SNAP$_i$-EGFP-cRaf: orange, K6-tag; purple, SNAP$_i$; green, EGFP; gray box, cRaf; yellow marker, EMCV IRES; brown, pac. (**j**) pPBbsr-ERK-KTR-mCherry: orange, K6-tag; gray box, ERK-KTR; red, mCherry; yellow marker, EMCV IRES; brown, blasticidin *S*-deaminase (bsr). (**k**) pPBpuro-K6-SNAP$_i$-EGFP-p85$_{iSH2}$: orange, K6-tag; purple, SNAP$_i$; green, EGFP; gray box, an iSH2 domain (residues 617–724) from human PI3K (p85α); yellow marker, EMCV IRES; brown, pac. (**l**) pCMV-mCherry-PH$_{Akt}$: red, mCherry; grey box, a PH domain (residues 1–148) from human Akt1. (**m**) pCMV-K6-SNAP$_i$-EGFP-Sos: orange, K6-tag; purple, SNAP$_i$; green, EGFP; gray box, Sos. (**n**) pPBbsr-MEK-P2A-mCherry-ERK: grey, FLAG-tag; purple, MEK1 from *Xenopus laevis*; black box, 2A self-cleaving peptide (P2A)[S12]; red, mCherry; blue, ERK2(K57R) from *Xenopus laevis*; yellow marker, EMCV IRES; brown, bsr.

### Supplementary Methods: Chemical Synthesis

#### General materials and methods

All chemical reagents and solvents were purchased from commercial suppliers (Watanabe Chemical Industries, Tokyo Chemical Industry, and Kanto Chemical) and used without further purification. Reverse-phase HPLC was performed on a Hitachi LaChrom Elite system with UV detection at 220 nm using a YMC-Pack ODS-A column (10 × 250 mm or 20 × 250 mm). $^{1}$H NMR spectra were recorded on a Bruker AVANCE III HD400SJ (400 MHz) spectrometer. $^{1}$H NMR chemical shifts were referenced to tetramethylsilane (0 ppm). High-resolution mass spectra were measured on a Thermo Scientific Extractive Plus Orbitrap mass spectrometer.

mgcBCP[S8] and compound **3** (BCP-COOH)[S13] were synthesized as described previously.

#### Reagent abbreviations

$Boc_2O$: di-*tert*-butyl dicarbonate
DIPEA: *N*,*N*-diisopropylethylamine
DMF: *N*,*N*-dimethylformamide
Fmoc-Adox-OH: Fmoc-8-amino-3,6-dioxaoctanoic acid
HBTU: *O*-(benzotriazole-1-yl)-*N*,*N*,*N'*,*N'*-tetramethyluronium hexafluorophosphate
HOBt: 1-hydroxybenzotriazole (monohydrate)
TFA: trifluoroacetic acid
TIPS: triisopropylsilane
TMS: tetramethylsilane

#### General methods for solid-phase synthesis

Compounds **1** (m$^{D}$cBCP) and **2** (adoxBCP) were synthesized manually on Sieber amide resin by standard Fmoc-based solid-phase peptide synthesis protocols. Fmoc deprotection was performed with 20% piperidine in DMF at room temperature for 30 min. Amino acid coupling reactions were performed at room temperature with a mixture of Fmoc-protected amino acid (3.1 eq.), HBTU (3.0 eq.), HOBt (3.0 eq.), and DIPEA (6.0 eq.) in DMF. All Fmoc deprotection and coupling steps were monitored by the Kaiser test.[S14] Unless otherwise stated, all washing procedures were performed with DMF.

### Synthesis of Compound 1

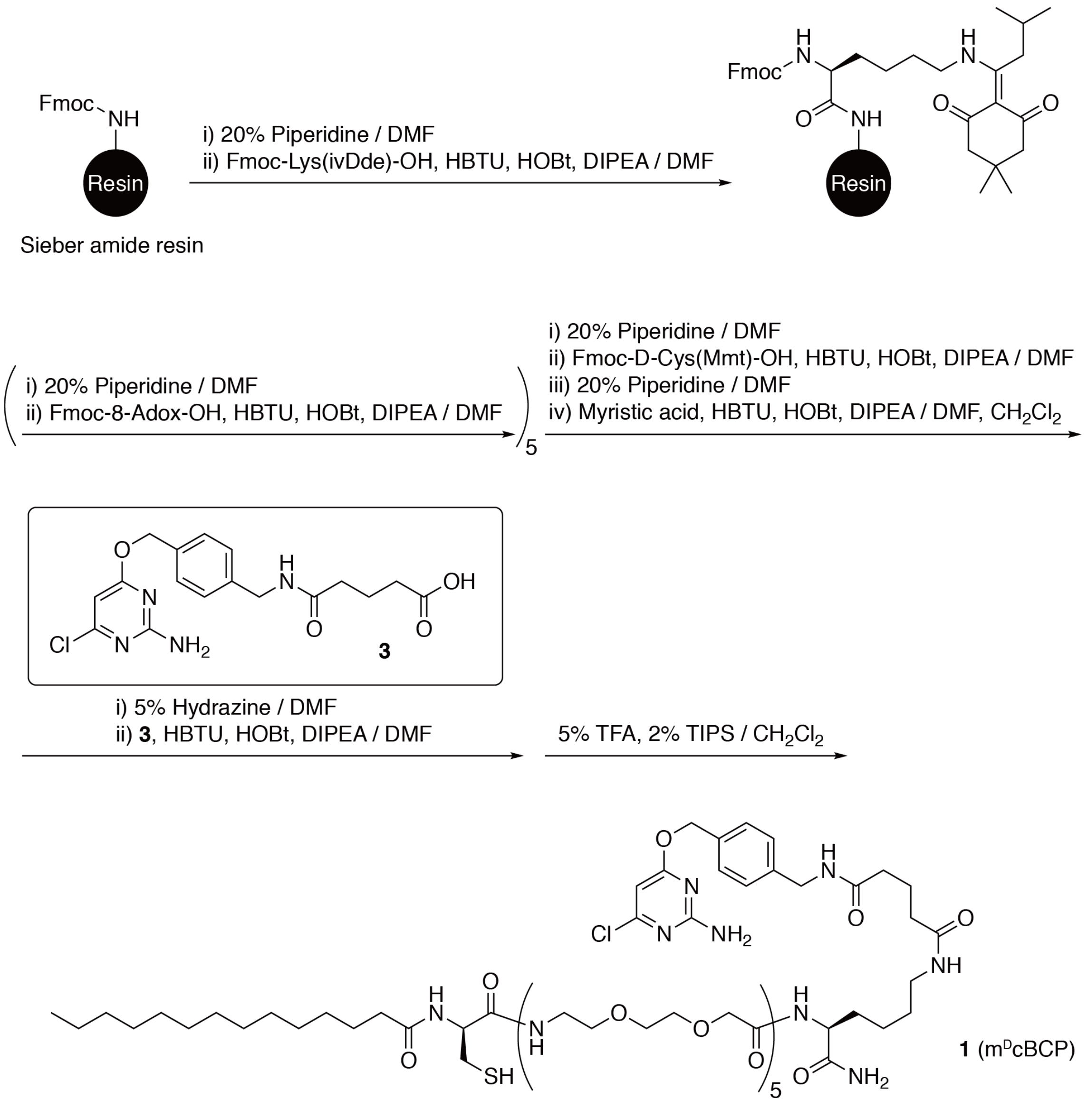


**Scheme S1.** Synthetic route of **1** (m$^{D}$cBCP)

Compound **1** (m$^{D}$cBCP) was synthesized manually on Sieber amide resin (0.69 mmol/g) (58 mg, 40 μmol). First, Fmoc-Lys(ivDde)-OH, Fmoc-Adox-OH (×5), and Fmoc-D-Cys(Mmt)-OH were coupled to the resin. The N-terminus was then myristoylated using a mixture of myristic acid (3.1 eq.), HBTU (3.0 eq.), HOBt (3.0 eq.), and DIPEA (6.0 eq.) in DMF/$CH_2Cl_2$ (1/1). After washing the resin with DMF, the ivDde group was deprotected by treatment with DMF containing 5% hydrazine monohydrate. The resin was then washed with DMF. **3** (BCP-COOH) was coupled to the side chain of the lysine with a mixture of **3** (3.1 eq.), HBTU (3.0 eq.), HOBt (3.0 eq.), and DIPEA (6.0 eq.) in DMF. After washing with DMF, MeOH, and $CH_2Cl_2$, the resin was dried *in vacuo*. Deprotection and cleavage from the resin was performed with $CH_2Cl_2$ containing 5% TFA and 2% TIPS. The crude product was precipitated by $Et_2O$ and purified by reversed-phase HPLC using a semi-preparative C18 column (a linear gradient of MeCN containing 0.1% TFA and 0.1% aqueous TFA) to afford **1** ($m^D$cBCP) as a white solid [15.7 mg, 23.6% (as a mono-TFA salt)].

Compound **1** ($m^D$cBCP)
$^1$H NMR (400 MHz, $CD_3OD$): δ 7.39 (d, *J* = 8.1 Hz, 2H), 7.29 (d, *J* = 7.9 Hz, 2H), 6.10 (s, 1H), 5.33 (s, 2H), 4.50–4.39 (m, 2H), 4.38–4.33 (m, 2H), 4.05–3.95 (m, 10H), 3.72–3.67 (m, 20H), 3.62–3.53 (m, 10H), 3.49–3.36 (m, 10H), 3.21–3.10 (m, 2H), 2.90–2.71 (m, 2H), 2.26 (t, *J* = 7.4 Hz, 4H), 2.21 (t, *J* = 7.4 Hz, 2H), 1.95–1.67 (m, 4H), 1.66–1.57 (m, 2H), 1.56–1.47 (m, 2H), 1.41–1.36 (m, 2H), 1.28 (s, 20H), 0.89 (t, *J* = 6.8 Hz, 3H).
HRMS (ESI): calculated for $[M+H]^+$, 1544.8047; found, 1544.8002.

### Synthesis of Compound 2

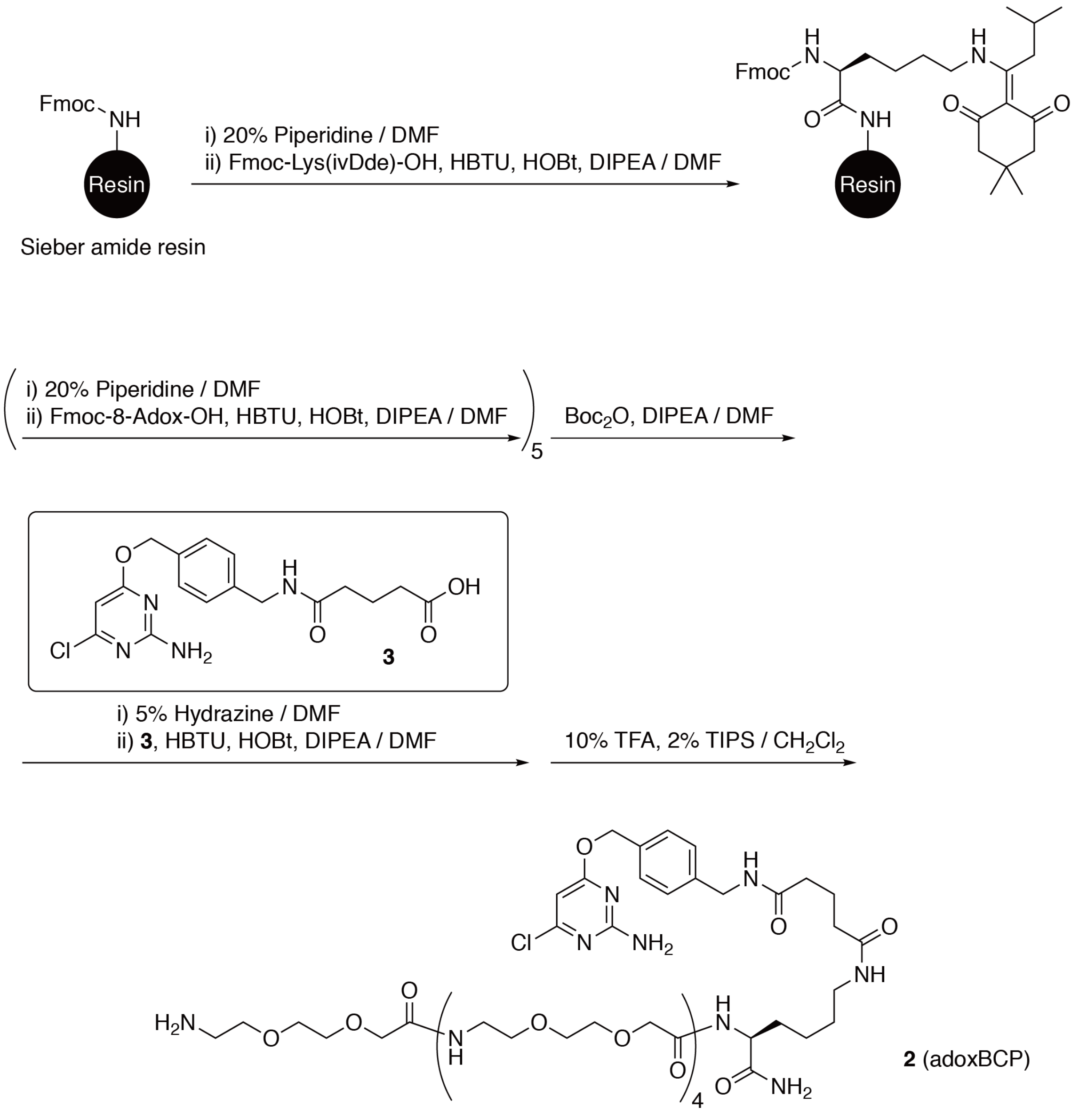


**Scheme S2.** Synthetic route of **2** (adoxBCP)

Compound **2** (adoxBCP) was synthesized on Sieber amide resin (0.69 mmol/g) (29 mg, 20 μmol). First, Fmoc-Lys(ivDde)-OH and Fmoc-Adox-OH (×5) were coupled to the resin. After Fmoc deprotection, the N-terminus was Boc-protected using a mixture of $Boc_2O$ (3.1 eq.) and DIPEA (6.0 eq.) in DMF. After washing the resin with DMF, the ivDde group was deprotected by treatment with DMF containing 5% hydrazine monohydrate. The resin was then washed with DMF. **3** was coupled to the side chain of the lysine with a mixture of **3** (3.1 eq.), HBTU (3.0 eq.), HOBt (3.0 eq.), and DIPEA (6.0 eq.) in DMF. After washing with DMF, MeOH, and $CH_2Cl_2$, the resin was dried *in vacuo*. Deprotection and cleavage from the resin was performed with $CH_2Cl_2$ containing 10% TFA and 2% TIPS. The crude product was precipitated by $Et_2O$ and purified by reversed-phase HPLC using a semi-preparative C18 column (a linear gradient of MeCN containing 0.1% TFA and 0.1% aqueous TFA) to afford **2** (adoxBCP) as a colorless oil [10.5 mg, 35.9% (as a di-TFA salt)].

Compound **2** (adoxBCP)
$^1$H NMR (400 MHz, $CD_3OD$): δ 7.38 (d, $J$ = 8.1 Hz, 2H), 7.28 (d, $J$ = 8.1 Hz, 2H), 6.10 (s, 1H), 5.33 (s, 2H), 4.46–4.39 (m, 1H), 4.36 (s, 2H), 4.00 (m, 10H), 3.72–3.63 (m, 22H), 3.61–3.55 (m, 8H), 3.48–3.42 (m, 8H), 3.18–3.11 (m, 4H), 2.26 (t, $J$ = 7.5 Hz, 2H), 2.20 (t, $J$ = 7.5 Hz, 2H), 1.90 (quin, $J$ = 7.5 Hz, 2H), 1.57–1.46 (m, 2H), 1.45–1.24 (m, 4H)
HRMS (ESI): calculated for $[M+H]^+$, 1231.5972; found, 1231.5924.

### Supplementary Methods: Molecular and Cell Biology Experiments

#### Plasmid construction

All of the cDNA and amino acid sequences of the constructs used in this study are listed in **Fig. S7** and **Fig. S8**. We used pPB-CAG.EBNXN (provided by Dr. Allan Bradley, Wellcome Trust Sanger Institute),[S15] pEGFP-C1 (Clontech), pmCherry-N1 (Clontech), and pmCherry-C1 (Clontech) as vector backbones. We also used pPBbsr[S8,S16] (blasticidin *S* resistance) and pPBpuro[S8] (puromycin resistance), as *piggyBac* donor vectors for the establishment of stable cell lines. All expression plasmids were generated using standard cloning procedures.

#### Cell culture and transfection

HeLa cells were obtained from the Cell Resource Center for Biomedical Research, Institute of Development, Aging and Cancer, Tohoku University. Cells were cultured in DMEM (Wako) supplemented with 10% heat-inactivated FBS (Biowest), penicillin (100 U/mL), and streptomycin (100 μg/mL) at 37°C under a humidified 5% $CO_2$ atmosphere. For transient expression experiments, cells were transfected using Lipofectamine LTX (Invitrogen) or 293fectin (Invitrogen) in accordance with the manufacturer's protocol.

#### Establishment of stable cell lines

A *piggyBac* transposon system[S15] was employed to establish HeLa cell lines stably expressing the indicated constructs. HeLa cells were cotransfected with a *piggyBac* donor vector (pPBpuro) encoding a desired protein(s) and pCMV-mPBase encoding the *piggyBac* transposase[S15,S17] (provided by Dr. Allan Bradley, Wellcome Trust Sanger Institute) using 293fectin (Invitrogen). Cells were selected with 2 μg/mL puromycin for at least 10 days. Bulk populations of selected cells were used.

#### Live cell imaging

Fluorescence imaging was performed with an IX83/FV3000 confocal laser-scanning microscope (Olympus) equipped with a PlanApo N 60×/1.42 NA oil objective (Olympus), a Z drift compensator system (IX3-ZDC2, Olympus), and a stage top incubator (Tokai Hit). Lasers used for excitation were as follows: 488 nm for EGFP, 561 nm for mCherry. Time-lapse live cell imaging was performed at 37°C. Fluorescence images were analyzed using the Fiji distribution of ImageJ.[S18] The cell outline was traced using the QuimP software.[S19]

**SLIPT assays**

To conduct the SLIPT assay, HeLa cells stably expressing K6-SNAP$_f$-Halo (established using pPBpuro-K6-SNAP$_f$-Halo), K6-SNAP$_i$-Halo (pPBpuro-K6-SNAP$_i$-Halo), K6-SNAP$_f$-ns-Halo (pPBpuro-K6-SNAP$_f$-ns-Halo), K6-SNAP$_f$-EGFP (pPBpuro-K6-SNAP$_f$-EGFP), or K6-SNAP$_i$-EGFP (pPBpuro-K6-SNAP$_i$-EGFP) were plated at $1.0\times10^5$ cells in 35 mm glass-bottomed dishes (Iwaki Glass) coated with collagen type I-C (Nitta gelatin). The cells were then cultured for 24 h at 37°C in 5% $CO_2$. For cells expressing HaloTag fusion proteins, HaloTag was labeled as described below before the SLIPT assay. The medium was changed to serum-free DMEM supplemented with penicillin (100 U/mL) and streptomycin (100 μg/mL) [DMEM(–)], and the cells were observed by time-lapse imaging before and after addition of the indicated compounds (10 μM) dissolved in DMSO (final DMSO concentration <0.1% v/v).

**HaloTag labeling**

Cells expressing HaloTag fusion proteins were stained as follows. After 24 h incubation of the cells at 37°C in 5% $CO_2$, the cells were washed with DMEM(–) and incubated with 50 nM HaloTag® TMR ligand (Promega) in DMEM(–) for 15 min at 37°C in 5% $CO_2$. The cells were then washed with DMEM(–) and incubated in (dye-free) DMEM(–) for 30 min at 37°C in 5% $CO_2$. The cells were washed with DMEM(–) once again and used for the SLIPT assay.

**HPLC analysis of mgcBCP degradation**

Cellular degradation of mgcBCP was analyzed as previously reported.[S10] Intact or K6-SNAP$_f$-Halo-expressing HeLa cells were plated at $4.0\times10^5$ cells in 60 mm plastic dishes (TPP) and cultured for 24 h at 37°C. The cells were washed with HEPES buffered saline (HBS) (25 mM HEPES, 119 mM NaCl, 5 mM KCl, 2 mM $MgCl_2$, 2 mM $CaCl_2$, 30 mM glucose, pH 7.4) (2 mL×2) and the medium was changed to 1 mL of HBS containing mgcBCP or m$^D$cBCP (10 μM). After incubation for 3 h at 37°C, the buffer solution was collected and filtered with a membrane filter (Millipore, Millex-LH, 0.45 μm). A total of 900 μL of the filtrate was analyzed by reversed-phase HPLC using a C18 column (YMC-

Pack ODS-A column, 10×250 mm) with a linear gradient of MeCN containing 0.1% TFA (A) and 0.1% aqueous TFA (B): A/B = 0/100 to 100/0 for 60 min. The HPLC profiles were monitored at 220 nm. The new product shown in **Fig. S1** was characterized by high-resolution mass spectrometry (Thermo Scientific Extractive Plus Orbitrap mass spectrometer). For control experiments, mgcBCP or $m^{D}$cBCP (10 μM) dissolved in HBS was incubated for 3 h at 37°C and analyzed by reversed-phase HPLC as described above.

**Synthetic Tiam1-mediated Rac1 activation and lamellipodia formation**

HeLa cells were plated at $0.2\times10^5$ cells in 35 mm glass-bottomed dishes coated with poly-L-lysine solution (Sigma). The cells were cotransfected with pCMV-Lifeact-mCherry and either pCMV-K6-$SNAP_f$-EGFP-Tiam1 or pCMV-K6-$SNAP_i$-EGFP-Tiam1. Twenty-four hours after transfection, the medium was changed to DMEM(–), followed by serum-starvation for 1 h. The cells were imaged before and after the addition of mgcBCP or $m^{D}$cBCP (10 μM).

**Synthetic cRaf-mediated ERK activation**

HeLa cells were plated at $1.0\times10^5$ cells in 35 mm glass-bottomed dishes coated with collagen type I-C. The cells were cotransfected with pPBpuro-K6-$SNAP_i$-EGFP-cRaf and pPBbsr-ERK-KTR-mCherry. Twenty-four hours after transfection, the medium was changed to DMEM(–), followed by serum-starvation for 1 h. The cells were imaged before and after the addition of $m^{D}$cBCP (10 μM).

**Synthetic PI3K-mediated PI(3,4,5)$P_3$ production**

HeLa cells were plated at $1.0\times10^5$ cells in 35 mm glass-bottomed dishes coated with collagen type I-C. The cells were cotransfected with pCMV-K6-$SNAP_i$-EGFP-iSH2 and pCMV-mCherry-$PH_{Akt}$. Twenty-four hours after transfection, the medium was changed to DMEM(–), followed by serum-starvation for 1 h. The cells were imaged before and after the addition of $m^{D}$cBCP (10 μM).

**Synthetic Sos-mediated Ras/ERK activation**

HeLa cells were plated at $1.0\times10^5$ cells in 35 mm glass-bottomed dishes coated with collagen type I-C. The cells were cotransfected with pPBpuro-K6-$SNAP_i$-EGFP-Sos and pPBbsr-MEK-P2A-mCherry-ERK. Twenty-four hours after transfection, the medium was changed to DMEM(–), followed by serum-starvation for 1 h. The cells were imaged before and after the addition of m$^{D}$cBCP (10 μM).

### References


S1. B. Mollwitz, E. Brunk, S. Schmitt, F. Pojer, M. Bannwarth, M. Schiltz, U. Rothlisberger and K. Johnsson, *Biochemistry*, 2012, **51**, 986–994.

S2. J. E. A. Wibley, A. E. Pegg and P. C. E. Moody, *Nucleic Acids Res*., 2000, **28**, 393–401.

S3. D. S. Daniels, C. D. Mol, A. S. Arvai, S. Kanugula, A. E. Pegg and J. A. Tainer, *EMBO J*., 2000, **19**, 1719–1730.

S4. T. D. Goddard, C. C. Huang, E. C. Meng, E. F. Pettersen, G. S. Couch, J. H. Morris and T. E. Ferrin, *Protein Sci.*, 2018, **27**, 14–25.

S5. G. Maryu, M. Matsuda and K. Aoki, *Cell Struct. Funct.*, 2016, **41**, 81–92.

S6. S. Regot, J. J.Hughey, B. T. Bajar, S. Carrasco and M. W. Covert, *Cell*, 2014, **157**, 1724–1734.

S7. B.-C. Suh, T. Inoue, T. Meyer and B. Hille, *Science*, 2006, **314**, 1454–1457.

S8. A. Nakamura, C. Oki, K. Kato, S. Fujinuma, G. Maryu, K. Kuwata, T. Yoshii, M. Matsuda, K. Aoki and S. Tsukiji, *ACS Chem. Biol.*, Article ASAP, DOI: 10.1021/acschembio.0c00024.

S9. L. J. Holsinger, D. M. Spencer, D. J. Austin, S. L. Schreiber and G. R. Crabtree, *Proc. Natl. Acad. Sci. U.S.A.*, 1995, **92**, 9810–9814.

S10. A. Nakamura, C. Oki, S. Sawada, T. Yoshii, K. Kuwata, A. K. Rudd, N. K. Devaraj, K. Noma and S. Tsukiji, *ACS Chem*. *Biol*., Article ASAP, DOI: 10.1021/acschembio.0c00014.

S11. Y. A. Bochkov and A. C. Palmenberg, *Biotechniques*, 2006, **41**, 283–288.

S12. J. H. Kim, S.-R. Lee, L.-H. Li, H.-J. Park, J.-H. Park, K. Y. Lee, M.-K. Kim, B. A. Shin and S.-Y. Choi, *PLoS One*, 2011, **6**, e18556.

S13. D. Srikun, A. E. Albers, C. I. Nam, A. T. Iavarone and C. J. Chang, *J. Am. Chem. Soc.*, 2010, **132**, 4455–4465.

S14. E. Kaiser, R. L. Colescott, C. D. Bossinger and P. I. Cook, *Anal. Biochem.*, 1970, **34**, 595–598.

S15. K. Yusa, R. Rad, J. Takeda and A. Bradley, *Nat. Methods*, 2009, **6**, 363–369.

S16. N. Komatsu, K. Aoki, M. Yamada, H. Yukinaga, Y. Fujita, Y. Kamioka and M. Matsuda, *Mol. Biol. Cell*, 2011, **22**, 4647–4656.

S17. J. Cadiñanos and A. Bradley, *Nucleic Acids Res.*, 2007, **35**, e87.

S18. J. Schindelin, I. Arganda-Carreras, E. Frise, V. Kaynig, M. Longair, T. Pietzsch, S. Preibisch, C. Rueden, S. Saalfeld, B. Schmid, J.-Y. Tinevez, D. J. White, V. Hartenstein, K. Eliceiri, P. Tomancak and A. Cardona, *Nat. Methods*, 2012, **9**, 676–682.

S19. P. Baniukiewicz, S. Collier and T. Bretschneider, *Bioinformatics*, 2018, **34**, 2695–2697.
